# Peering into the world of wild passerines with 3D-SOCS: synchronized video capture and posture estimation

**DOI:** 10.1101/2024.06.30.601375

**Authors:** Michael Chimento, Alex Hoi Hang Chan, Lucy M. Aplin, Fumihiro Kano

## Abstract

1. Collection of large behavioural data-sets on wild animals in natural habitats is vital in ecology and evolution studies. Recent progress in machine learning and computer vision, combined with inexpensive microcomputers, have unlocked a new frontier of fine-scale markerless measurements.
2. Here, we leverage these advancements to develop a 3D Synchronized Outdoor Camera System (3D-SOCS): an inexpensive, mobile and automated method for collecting behavioural data on wild animals using synchronized video frames from Raspberry Pi controlled cameras. Accuracy tests demonstrate 3D-SOCS’ markerless tracking can estimate postures with a 3mm tolerance.
3. To illustrate its research potential, we place 3D-SOCS in the field and conduct a stimulus presentation experiment. We estimate 3D postures and trajectories for multiple individuals of different bird species, and use this data to characterize the visual field configuration of wild great tits (*Parus major*), a model species in behavioural ecology. We find their optic axes at approximately ±60° azimuth and −5° elevation. Furthermore, birds exhibit functional lateralization in their use of the right eye with conspecific stimulus, and show individual differences in lateralization. We also show that birds’ convex hulls predicts body weight, highlighting 3D-SOCS’ potential for non-invasive population monitoring.
4. 3D-SOCS is a first-of-its-kind camera system for wild research, presenting exciting potential to measure fine-scaled behaviour and morphology in wild birds.

## 1 Introduction

Automated, large-scale data collection methods to measure animals’ behaviour in their natural habitat are of growing importance for understanding social systems (Farine et al. 2014; Farine et al. 2015b), migration and dispersal (Flack et al. 2018; Klarevas-Irby et al. 2021; Flack et al. 2022), disease and information transmission (Aplin et al. 2015; Farine et al. 2015a; Fang et al. 2020) responses to climate and weather events (Hawkes et al. 2009; Nourani et al. 2023), and conservation impacts (Lahoz-Monfort et al. 2021; Tuia et al. 2022; Couzin et al. 2023). Increasingly, behaviour in the wild is measured using animal-borne sensor technologies. For example, GPS or accelerometers are employed to measure large scale movement patterns and fine-scale behaviours (Wilmers et al. 2015), but are often limited by the size, battery life and data storage of the device (T. A. Wild et al. 2023). To follow smaller animals or to increase the scope of studies, Passive Integrative Transponder (PIT) tags that utilize Radio Frequency Identification (RFID) technology are commonly used. Thanks to its small size and durability, RFID technology is widely employed to monitor wild bird and rodent populations (Bonter et al. 2011; König et al. 2015; Iserbyt et al. 2018; Raulo et al. 2024). Analyzing co-occurrence data from RFID tags has provided insights into social interactions, such as parental provisioning (Maldonado-Chaparro et al. 2021) and social networks (Aplin et al. 2013; Farine et al. 2014; Beck et al. 2020). Additionally, the widespread adoption of low-cost single-board computers, such as Raspberry Pi (Jolles 2021), has further enabled the use of RFID technology in experimental devices designed to study cognition (Aplin et al. 2015; Cauchoix et al. 2022; S. Wild et al. 2023; Chimento et al. 2024), or to manipulate social associations in wild populations (Kings et al. 2023; Beck et al. 2024).

While RFID technology provides information about the presence and absence of individuals near RFID antennas, it does not provide detailed information about their behaviours. Furthermore, RFID readings may be affected by a variety of environmental conditions such as humidity. An emerging method that has the potential to address this limitation involves the use of computer vision and machine learning tools (A. Mathis et al. 2020; Mackenzie Weygandt Mathis et al. 2020; Luxem et al. 2023), a development facilitated by a concomitant increase in the volume and resolution of image data (Griebling et al. 2022; Ehlman et al. 2023). Computer vision enables the automatic quantification of fine-scaled behaviours from images, not only increasing the volume of behavioural data but also enhancing the objectivity of behaviour coding. Although initial applications were primarily in controlled laboratory settings (Alarcón-Nieto et al. 2018; A. Mathis et al. 2018; Dunn et al. 2021; Karashchuk et al. 2021; Walter et al. 2021; Pereira et al. 2022; Nagy et al. 2023; Naik et al. 2023), computer vision is also being applied in the field. It has been used to monitor animals from drones (Graving et al. 2019; Koger et al. 2023; Schad et al. 2023), quantify courtship displays (Janisch et al. 2021), analyze parental care behaviours (Chan et al. 2023), study underwater interactions (Dunkley et al. 2023), identify individuals (Ferreira et al. 2020; Schofield et al. 2023), and more.

A particular branch of computer vision that is revolutionizing the way we measure animal behaviour is 2D and 3D posture estimation. Posture estimation refers to the task of estimating the position of morphological keypoints (e.g eyes, limbs, joints) from an image. With the recent development of frameworks like DeepLab-Cut, DeepPoseKit and SLEAP, multi-animal posture estimation in 2D and subsequent automated behavioural analysis have become possible (A. Mathis et al. 2018; Graving et al. 2019; Lauer et al. 2022; Pereira et al. 2022). In 3D, frameworks like Anipose, DANNCE and 3D-MuPPET (Dunn et al. 2021; Karashchuk et al. 2021; Waldmann et al. 2024) have also been developed to estimate postures in 3D using multiple camera views (but see Gosztolai et al. 2021 for single view 3D estimation). However, these frameworks are often limited to tracking a single animal, and only a few recent studies have demonstrated the ability to estimate the 3D postures of multiple individuals simultaneously (see An et al. 2023; Y. Han et al. 2024; Waldmann et al. 2024). Furthermore, while applications of 2D posture estimation in the wild is widespread (e.g. Graving et al. 2019; Labuguen et al. 2021; A. Mathis et al. 2021; Wiltshire et al. 2023; Ye et al. 2024), applications of 3D postures in the wild is limited, with notable exceptions including posture estimation of a single cheetah (Joska et al. 2021) and of three captive pigeons in an outdoor garden (Waldmann et al. 2024). The lack of adoption of 3D animal posture tracking can be attributed to a few main challenges. Firstly, algorithms for multiple individuals are less mature due to correspondence matching (identifying and matching the same points across different frames or views) and object trajectory tracking (Xiao et al. 2023). Secondly, 3D tracking often requires multiple synchronized camera views, which is either expensive or difficult to set up outside of the lab (Janisch et al. 2021).

Yet, the potential benefits of applying 2D and 3D posture tracking in the wild are immense. Automatic posture estimation is not only less labor-intensive but also achieves greater objectivity than manual behaviour coding based on ethograms (Anderson et al. 2014). Even though recent benchmark datasets like KABR (Kholiavchenko et al. 2024), MammalNet (J. Chen et al. 2023) and PanAf20K (Brookes et al. 2024), and tools like tools like LabGym (Hu et al. 2023) or YOLO-Behaviour (Chan et al. 2024) is advancing behavioural classification directly from videos, posture based methods often provide finer scale information for behavioural kinematics. For example, classification of posture-based measurements can provide direct information about individual and group behaviours such as social and dominance interactions (Xiao et al. 2023; Y. Han et al. 2024), decision making and information transfer (Koger et al. 2023; Delacoux et al. 2024) and courtship behaviour (Janisch et al. 2021; Mitoyen et al. 2021). Marker-based systems have been recently employed to measure the 3D head orientation of birds, presenting an effective method for estimating visual field use, lateralization of eye use, and attentional foci of birds (Itahara et al. 2022; Kano et al. 2022; Nagy et al. 2023; Delacoux et al. 2024; Itahara et al. 2024). However, marker-based systems are limited to the laboratory, highlighting the need for the development of markerless systems that could extend this approach to field setups.

Here, we introduce 3D synchronized outdoor camera system (3D-SOCS), a mobile, flexible and low-cost system that combines computer vision and sensor technology to measure proximity, identities and postures of wild animals (Figure 1). Our system is unique compared to published frameworks, integrating multi-animal posture estimation, trajectory tracking and individual identification, as well as providing hardware and software guidelines for deployment (Table 1). An additional novelty was our custom camera encoder software for frame synchronization between RaspberryPis, which could be used in any application requiring frame synchronized videos.

**Table 1:** Contemporary studies of markerless 3D animal tracking. Columns include reference, species tracked, whether tracking was done in the wild or captivity, whether multiple individuals were tracked simultaneously, whether trajectory, pose were tracked, whether re-identification was possible after individuals left the frame, and whether the study presented software or hardware. 2D tracking studies have been excluded.

| Study | Species | Conditions | Multiple | Trajectory | Posture | Re-ID | Focus |
| --- | --- | --- | --- | --- | --- | --- | --- |
| *Anipose*<br>Karashchuk et al. 2021 | fruit fly<br>mouse<br>human | captive<br>(indoor) | n | n | y | n | software |
| *DANNCE*<br>Dunn et al. 2021 | rat<br>mouse<br>marmoset<br>chickadee | captive<br>(indoor) | n | n | y | n | software |
| *Liftpose3D*<br>Gosztolai et al. 2021 | fruit fly<br>mouse<br>rat<br>macaque | captive<br>(indoor) | n | y | y | n | software |
| *Deepfly3D*<br>Günel et al. 2019 | fruit fly | captive<br>(indoor) | n | n | y | n | software |
| *MAMMAL*<br>An et al. 2023 | pig<br>dog<br>mouse | captive<br>(indoor) | y | y | y | n | software |
| *SBeA*<br>Y. Han et al. 2024 | rat<br>parrot | captive<br>(indoor) | y | y | y | n | software |
| *OpenMonkeyStudio*<br>Bala et al. 2020 | macaques | captive<br>(indoor) | y | y | y | n | software |
| Badger et al. 2020 | brown-headed<br>cowbird | captive<br>(outdoor<br>aviary) | n | n | y | n | software |
| Xiao et al. 2023 | brown-headed<br>cowbird | captive<br>(outdoor<br>aviary) | y | y | n | y | software |
| *3D-MuPPET*<br>Waldmann et al. 2024 | pigeon | captive<br>(outdoor) | y | y | y | n | software |
| Ling et al. 2018 | jackdaw<br>rook | wild<br>(outdoor) | y | y | n | n | hardware<br>software |
| Janisch et al. 2021 | collared<br>manakin | wild<br>(outdoor) | n | y | n | n | hardware<br>software |
| *AcinoseT*<br>Joska et al. 2021 | cheetah | wild<br>(outdoor) | n | y | y | n | software |
| *3D-SOCS* (ours) | *great tit*<br>*blue tit* | *wild*<br>*(outdoor)* | y | y | y | y | *hardware*<br>*software* |

**Figure 1:**
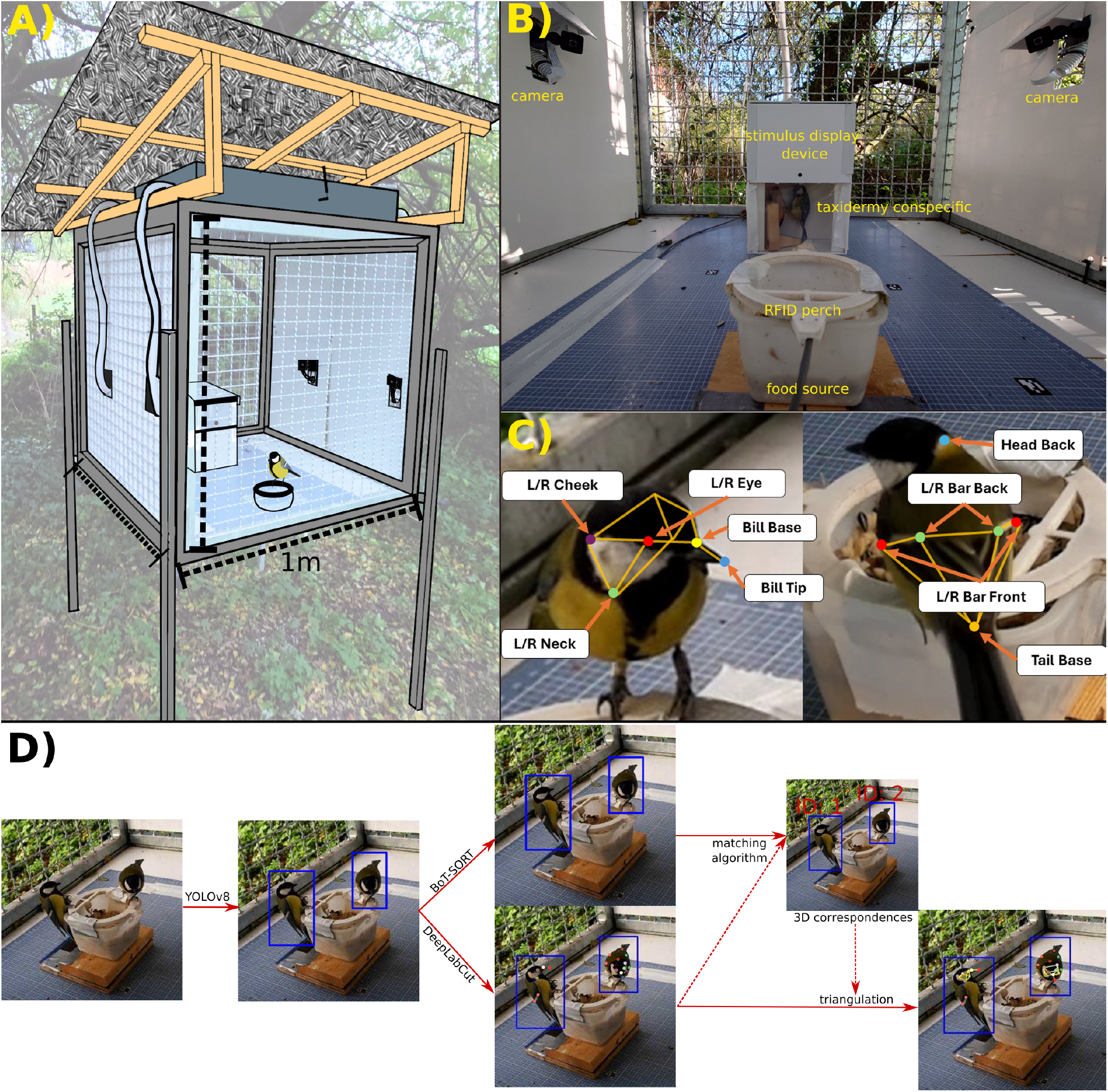
Overview of 3D-SOCS. A) Our mobile data collection system was housed in an elevated 1×1m cage with a roof for protection from the elements. The cage was designed to exclude large non-target species but to allow easy access for great tits. We installed white PVC plate walls to remove possible distractions in their lateral visual field. B) Whenever a tagged bird landed on the feeding perch antenna (foreground), the PLDC glass of the stimulus presentation device (background) turned transparent after a 1s delay, revealing the stimulus. Six frame-synchronized cameras placed around the cage filmed birds from various angles as they regarded the stimulus. C) We extracted keypoints on the head and body of birds from each frame, and used this information to create a 3D representation of the bird. D) Summary of the 3D tracking pipeline applied to videos.

We used 3D-SOCS to track the 3D posture of individuals from a population of great tits (*Parus major*) and blue tits (*Cyanistes caeruleus*) within 3mm accuracy. To showcase the capabilities of 3D-SOCS, we conducted a stimulus presentation experiment to examine the use of visual fields and lateralization of eye use in great tits, based on the tracking of their head orientation in 3D (Itahara et al. 2022; Kano et al. 2022; Itahara et al. 2024). While our system typically uses six cameras, we demonstrate that 3D-SOCS can operate effectively with just two cameras as a minimal setup. We also provide detailed documentation and code to enable other researchers to deploy the system (see section 5).

## 2 Material and Methods

### 2.1 Study species and site

Great tits (Parus major) are a common passerine songbird with a generalist diet found in woodlands and gardens throughout Europe and Asia (Perrins 1979). They are an established study species in behavioural ecology (Gibb 1950), with extensive previous study of their breeding, foraging and social behaviour (Fisher et al. 1949; Lefebvre 1995; Estók et al. 2010; Morand-Ferron et al. 2011; Aplin et al. 2015; Chimento et al. 2021).

The study took place in the woodland surroundings of the Max Planck Institute of Animal Behavior, located in Möggingen, Germany (47.765088, 8.997779). Wild tits were caught using mist-nets within a 1 km radius of the institute. Once caught, birds were ringed and fitted with a passive transponder (PIT tag) under ethical approval from the Regierungspräsidium Freiburg (35-9185.81/G-22/070) with rings provided by the Vogelwarte Radolfzell. While ringing, we also recorded wing and tarsus length, weight, age class (juvenile *<* 1 yr, adult ≥ 1 yr), and sex based on plumage.

### 2.2 3D-SOCS

3D Synchronized Outdoor Camera System (3D-SOCS) is a versatile markerless 3D posture tracking system for small passerine birds that runs on a battery and can be readily deployed in the field (Figure 1A). We designed the system to be flexible, in terms of the number of cameras it uses, where it can be deployed (battery powered), and the type of experimental apparatus that it be deployed with. It is also affordable compared to proprietary alternatives (6 camera setup €2500; minimal setup €1700), as the system is based on relatively cheap micro-computers. See supplementary methods section S1.1 for detailed description of the hardware and software required to deploy and collect data with 3D-SOCS.

While 3D-SOCS can use any rigid frame to mount the cameras, we used a 1×1m metal cage that housed our cameras and experimental devices (Figure 1A). The cage mesh was large enough to let small passerine birds in and out, but protected objects inside of the cage from other non-target visitors. We covered the left, right and bottom of the cage with white PVC plate to protect to prevent visual distractions from outside of the cage. The system was powered by a rechargeable battery unit.

Within the cage, we mounted six frame-synchronized cameras at various angles focused on a single feeder where birds were expected to land, consisting of a small bowl filled with a sunflower seed and peanut mixture (Figure 1B). Each camera was controlled by a RaspberryPi 4 Compute Module (CM4) micro-computer using PiCamera2 python library. A seventh CM4 was designated as the lead, and it acquired time from GPS satellites using a Timebeat GPS/GNSS module. The other six CM4 synchronized their hardware clocks to this time using Precision Time Protocol (PTP). Our custom encoder disciplined cameras framerates to the synchronized system clock of its CM4, achieving frame captures on each camera with sub-millisecond differences at 30 fps. Recording was triggered by a computer vision based motion-detection algorithm, which also ran on the lead CM4. When motion was detected, it triggered the other six cameras to begin recording using a GPIO signal. Recordings lasted for 60 seconds each. Further details of the synchronized video capture are available in Text S1.1, Figure S1, and online in our Github repository (see section 5).

Utilizing these synchronized cameras, we could reconstruct the 3D postures of birds visiting the feeder, and we were able to track multiple birds at the same time. Our target species was great tits, although a smaller number of tagged blue tits were also present, so we extended the model to detect their keypoints. We adopted a top-down approach similar to Waldmann et al. 2024 by: 1) localizing individual birds in each frame using YOLOv8 (Jocher et al. 2023), 2) estimating 14 unique 2D keypoints (Figure 1C) with DeepLabCut (A. Mathis et al. 2018; Kane et al. 2020), and 3) triangulating 2D postures into 3D (Figure 1D). For each unique individual recorded in the study (*N* = 57 great tits, *N* = 20 blue tits), we collated all 3D information and calculated the median head and body skeleton, then ensured the detections fit the median head skeletons by computing the estimated rotation translation of the skeleton in 3D space. This approach leverages the assumption that certain keypoints on the head of a bird form a rigid body (Naik et al. 2023), allowing for missing keypoints to be filled and relative distances between keypoints to be fixed, leading to more consistent detections. We refer to section 2.6 for the detailed 3D tracking pipeline and supplementary video S1 for qualitative results.

### 2.3 Stimulus presentation experiment

In order to test and showcase the capabilities of 3D-SOCS, we did a stimulus presentation experiment with the aim of estimating the visual field use of great tits. We began with a 7 day habituation phase, where we placed a large plate of seeds and mealworms in the arena to habituate birds to 3D-SOCS. We also used this period to troubleshoot the data collection pipeline.

For 5 days (November 5th to November 10th, 2023) following the habituation period, we deployed a stimulus presentation box and a feeder with an RFID antenna, which could read unique identities for birds wearing a passive-integrated transponder (PIT) leg band (Figure 1). When a PIT-tagged bird landed on the reader, our stimulus presentation device revealed the stimulus (see following section for details). We measured the head rotation angles relative to the stimulus within 3 seconds of the stimulus presentation to determine the areas of the visual fields most frequently used by birds. This area should encompass the optic axis they use, which is situated close to the birds’ fovea (area *centralis*). The width of this area is likely influenced by the degree of eye movement as well as by measurement errors (Pratt 1982; Bischof 1988; Butler et al. 2018; Kano et al. 2022).

We tested birds in four conditions presenting four different stimuli: 1) No stimulus condition as control for the clicking sound of the switchable glass turning on, 2) food stimulus condition presenting a small dish filled with mealworms, a high-reward food item, 3) food video stimulus condition presenting a looped animation of a single mealworm that wiggled in place on a screen (Huawei MediaPad T2-8 Pro, 60 Hz 8” IPS screen with 1200×1920px resolution), and 4) social stimulus condition presenting a taxidermy female adult great tit. Using a taxidermy bird allowed us to control the identity, posture and location of the observed bird. In total, we estimated head posture data of 34, 587 frames from *N* = 57 great tits (26 females, 29 juveniles). Each bird was recorded as visiting 3D-SOCS on average 18 times (range: 1-62). The orienting responses to the visual stimulus were defined as moments when the bird’s head was stationary (head-fixation); frames in which the bird exhibited rapid saccade-like head movements were excluded from the analysis.

### 2.4 Stimulus presentation device

In order to standardize the stimulus presentation to individuals, we constructed a small box which held a stimulus behind a piece of polymer dispersed liquid crystal (PDLC) film, a material that is in an opaque or transparent state depending on whether a current is supplied. The device was controlled by a RaspberryPi 4 microcomputer with custom Python software. The software monitored a perch equipped with an energized antenna that was placed directly over the food source. When a PIT tagged bird landed on the perch, an RFID reader (PriorityOne brand) read the individual identity of the bird. After a delay of 1s, the PDLC film then became transparent, revealing the stimulus. It remained transparent until the bird was detected as leaving the perch. The device recorded the bird’s identity and a timestamp of when the bird arrived and left.

### 2.5 Datasets and model training

We trained two separate models, YOLOv8-large (Jocher et al. 2023), an object detection model to localize a bird in frame, and DeepLabCut (DLC; (A. Mathis et al. 2018)) with pre-trained ResNet50 as the 2D posture estimation model. For object detection, we manually annotated a dataset with 1601 frames randomly sampled across the views, with bounding boxes labelled “Great tit” and “Blue tit”. Out of these images, we further annotated 905 bird instances with 14 unique keypoints if it was visible. These keypoints included: bill tip, bill base, L/R eye, L/R neck, L/R cheek, head back, L/R wing bar front, L/R wing bar back and tail base (Figure 1C). We used default augmentation parameters for YOLO and custom image augmentation and learning rate for DLC. All annotations were done using the open source software Label Studio (Tkachenko et al. 2024). The complete keypoint and bounding box dataset is available (see section 5).

### 2.6 3D posture estimation

We briefly describe the processing pipeline, although full details can be found in section S1.2. We used a laser-printed ChArUco checkerboard generated using the opencv library for calibration of camera intrinsics and extrinsics at the beginning of each day of data collection. Intrinsic calibration refers to each camera’s internal parameters (e.g. focal length, lens distortion), and extrinsic calibration refers to parameters that describe the relative positions between each camera. Daily calibration was required due to drift introduced by slight movements in cameras over the day, and temperature fluctuations that can affect lens parameters of cameras (Table S1). Figure 1D illustrates the full tracking pipeline for 3D posture estimation. We first detected bounding boxes of all birds present in each frame using YOLOv8 and then tracked birds across frames in 2D using BoT-SORT to provide a unique identity for each bird (Aharon et al. 2022). Bounding boxes were used to crop frames of birds, which were then fed into a single animal DeepLabCut model (A. Mathis et al. 2018), to generate 2D keypoint predictions for each bird per view. Next, we estimated correspondence between individuals across views for every frame through iterative matching, based on Huang et al. (C. Huang et al. 2020), then matched 2D tracks by pooling all matches and grouping tracks above a certain threshold of match rate. We used the matched correspondences to triangulate 2D detections into 3D posture estimates per frame using bundle adjustment by minimizing the reprojection error of a predicted 3D point across all cameras.

We implemented several post-processing steps, including interpolation and smoothing. We filtered outlier 3D keypoints when the reprojections were outside of any camera frame. We removed keypoints that fell beyond a set boundary of the head or body, defined by 1 standard deviation away from the mean position of the overall object. Finally, since head keypoints were rigid relative to each other for each bird, we calculated an average “skull” for each individual by taking the median measurements for each point in the head coordinate system. If at least 3 points of a bird’s skull were detected, we could use the average skull to infer the position of any missing keypoints by computing its rotation and translation in space. We leveraged this when testing stereo cameras as a minimal set-up for 3D-SOCS. Stereo cameras are inherently more challenging since the number of points detected by both cameras is limited. However, as long as 3 points were detected, we used a species-specific median skeleton to fill in missing keypoints, providing 3D keypoint estimates of the whole bird.

### 2.7 System accuracy test

To validate the accuracy of 3D-SOCS, we carried out two system accuracy tests. Firstly, we randomly sampled 113 frames and manually annotated all keypoints in 2D from all 6 views (678 2D frames). These annotations were then triangulated with the same pipeline to get 3D ground truth. However, this evaluation is incomplete, since systematic errors like calibration or synchronization errors will still influence the ground truth data in the same way.

In order to thoroughly evaluate the 3D tracking system, we performed a second accuracy test by deploying the system with the same camera configuration in a highly accurate marker-based motion capture system SMART-BARN (Nagy et al. 2023). The goal of the test was to determine the mean and variance of the absolute distance error and rotational error of a given object for 3D-SOCS, against independent estimates from SMART-BARN. To inform future researchers on the trade-off between camera settings, we also tested two camera resolutions: a narrow field of view (1920×1080px) that is focused on the food tray, and a wide field of view (1640×1280px) which can capture a wider area of the 1 cubic-meter platform.

In brief, a taxidermy great tit and an ArUco QR code were systematically rotated and moved around the volume for approximately 20 minutes per camera resolution. Since the environment and taxidermy great tit looks slightly different from the data collected by 3D-SOCS in the field, we also fine-tuned the DeepLabCut and YOLO model using 100 additional annotated frames of the taxidermy tit, to provide the most ideal conditions to evaluate the accuracy of the system. We refer to section S1.4 for a detailed description of the workflow.

For both system accuracy tests, we measured the absolute distance and rotation error of a set of head keypoints (bill tip, L/R eyes), to determine the accuracy of the system for measuring visual fields of the birds. In addition, we also tested for the number of cameras required by iteratively removing the number of cameras in the system, and calculating the same accuracy metrics.

### 2.8 Species classification and trajectory tracking validation

Finally, we evaluated the species classification and trajectory tracking performance in 3D-SOCS. To evaluate the performance of species classification, we used the validation dataset consists of annotations of blue tit and great tit bounding boxes (N = 228), to compare the fine-tuned YOLOv8 model trained for 3D-SOCS with off-the-shelf methods. For off the shelf methods, we used the same YOLOv8-large model, but pre-trained on the COCO dataset (Lin et al. 2014), which includes the “bird” class. Detected birds were then inputted into the zero-shot BioCLIP model (Stevens et al. 2024), a foundation model for species identification. Since the pre-trained YOLO model consistently missed birds in the frame, we also directly cropped images of birds from the ground truth bounding boxes as input into zero-shot BioCLIP, to evaluate species identification performance.

Next, we evaluated the trajectory tracking performance in 3D-SOCS. From our dataset, we identified trials where multiple birds were simultaneously tracked for more than a third of the trial length, then visualized the trials by overlaying tracking ID on the birds (N = 111). We then manually reviewed each video and categorized videos as 1) all individuals correctly tracked, 2) same ID assigned to different birds, or 3) different IDs assigned to the same bird, with 2 and 3 being forms of ID switches. We then computed overall tracking accuracy across these trials, in terms of the proportion of trials that had no ID switches.

### 2.9 Field of view and lateralization analysis

Assuming that birds were most likely to regard the stimulus immediately after its display, we limited the posture data to include only the 3 seconds following the stimulus onset. For each frame, we transformed the 3D coordinate of the stimulus to the local head coordinate system, then calculated the position of the stimulus using a modified spherical coordinate system, where azimuth and elevation of 0 corresponded to the forward vector between the mid-point of the eyes and the bill tip (Figure S2).

We used a set of filters to subset frames where a bird was likely attending to the stimulus. Birds use saccadic head movements, characterized by a rapid movement of the head followed by a longer fixation (Kano et al. 2018). It should be noted that due to the limited temporal resolution of our RGB cameras (30 fps), our system was unable to detect small and brief head-saccades. Our aim was to extract head fixations that were separated by relatively large saccade-like movements, rather than to detect the head-saccades themselves. We first computed the angular change of a bird’s head between each frame and defined saccades as changes of *>* 500° per second (16.7°per frame). We then categorized frames between these detected saccades as fixation frames. Since the detected saccades that were very short or very long could potentially be attributed to noise in 3D posture estimation, we disregarded those saccades that were shorter than two frames (*<* 66*ms*) or longer than five frames (*>* 165*ms*). We also disregarded saccade amplitudes of *<* 33°, manually extracted from Figure S3, and fixation periods of 1 frame or less. To exclude frames where birds were eating seeds with their heads down (likely focusing on the food), we filtered out frames in which the bill tip was within a defined cuboid around the food tray.

We fit a Bayesian multivariate mixture model to this data to estimate the optic axes. Each observation was a two-dimensional vector representing the head azimuth and elevation relative to the stimulus. The likelihood of an observation was modeled as a mixture of *K* = 2 two-dimensional Gaussians (representing the lateral fields of their left and right eyes) with each component’s mean vector composed of an azimuth and elevation component (*µ*_az.,*s*_[*k*], *µ*_elev._) and scale vector (*σ*_az._[*k*], *σ*_elev._). Parameters and priors are summarized in Table S2. Azimuth means, standard deviations, and mixing proportions were estimated for each stimulus, but the elevation was estimated via complete pooling.

As this model was intended to estimate the head elevation and optic axes and could not account for varying effects of individual ID, we fit a secondary Bayesian logistic model with varying effects. We subset data to measurements of head elevation between −15° and 5° (informed by the first model and rotation error from the system accuracy tests) to conservatively only include observations of birds likely attending to stimuli. We coded a binary variable representing when a bird favored their right lateral field as 1 when azimuth was between 50° and 70°, and 0 when azimuth was −50° and −70°. This variable was modelled as a Bernoulli distribution with probability *θ*, composed of a sum of varying intercepts for stimulus type and individual ID.

### 2.10 Estimation of body surface area as proxy for weight

Using body keypoints (L/R wing bar front, L/R wing bar back and tail base), we constructed a median body skeleton for each individual bird across all trials, then defined a sparse mesh using the trimesh python library (Dawson-Haggerty et al. 2019). From this, we computed the surface area of a 3D convex hull as a proxy for body size. To test the relationship between the convex hull and weight in great tits, we used a Bayesian linear model where scaled weight was modeled as a normal distribution whose mean was predicted by scaled surface area and a varying intercept for species class. Each bird’s weight was calculated as the mean of all its measured weights across captures from our catalogue of ringing data, excluding its weight as a nestling.

### 2.11 Bayesian model fitting

All models described above were fit using Hamiltonian Markov Chain Monte Carlo (MCMC). Models were run using 5 chains and a minimum of 5000 iterations, with 2500 warm-up iterations, with all estimates based on over at least 1976 effective samples from the posterior (range: 1976–18955). All models were fitted using R v. 4.3.2 with Stan v. 2.32.2 (R Core Team 2018; Stan Development Team 2021). Good model convergence was confirmed from evaluating rank histograms and ensuring parameters had Gelman–Rubin’s statistic ≤ 1.01 (Vehtari et al. 2021). Given the complexity of the Gaussian mixture model used to estimate the optic axes, we further validated this model using simulated data (see section 5).

## 3 Results

### 3.1 System accuracy

We first evaluated the synchronization performance of 3D-SOCS, by comparing the time offset for the frame capture time for each camera compared to the mean time across all 6 cameras. Frames were highly synchronized, with a mean offset of 1.15 × 10^*−*5^*ms* and 99% intervals of -22*ms* − 28*ms*, within the frame-by-frame difference of 33*ms* at 30fps. We then evaluated 3D-SOCS to ensure all measured keypoint estimates of wild birds were accurate throughout the experimental period. Overall, we achieved low median 3D error, 2D detection error and 3D reprojection error of 1.95mm, 0.24px and 4.61px respectively, and an average of 18% frame loss rate across all keypoints (Tables S3, S4, S5). Body keypoints were lost more frequently than head keypoints (Head: 13%, Body: 28%). We also explored how differing numbers of cameras affected accuracy of detection based on multi-view manual annotations (Table S3, Figure S4). Incrementally reducing the number of cameras from six to two cameras all yielded similarly accurate results, but with a higher variance and frame loss rate as the number of cameras decreased.

We further validated the overall accuracy of the 3D-SOCS by comparing 3D-SOCS’ estimates with estimates from a marker-based motion capture system with sub-millimeter precision (SMART-BARN (Nagy et al. 2023)), which we treated as the ground truth. 3D-SOCS’ measurements for head keypoint locations were normally distributed around 0, for both the complete and stereo setup (visualized in Figure S5, S6). The upper 95% highest density interval (HDI) of keypoint detections were no further than 3mm from their ground-truth position, with a median deviation of 1.05mm (Table S6, S7, S8). Distributions of absolute rotation errors for yaw, pitch, and roll were also overall distributed around 0, with an upper 95% HDI between 7° and 11° depending on the rotation type (Table 2). We tested two different camera resolutions corresponding to narrow (1920×1080px) and wide (1640×1280px) fields of view. While 1080p resolution resulted in slightly better estimations of yaw and pitch, the two resolutions were generally comparable. In summary, our findings show no evidence of systematic deviations from the ground-truth values in the measurements.

**Table 2:** Summary of system accuracy tests with six cameras. Median absolute errors and upper 95% highest density interval (HDI) in parenthesis. Values of head keypoint positions and roll, pitch, yaw angles measured by 3D-SOCS were compared to SMART-BARN as ground truth. A value of zero would indicate a measurement that matched ground truth. Keypoints estimated from six cameras, recorded at either a resolution of 1920×1080px (narrow FOV) or 1640×1280px (wide FOV).

| Resolution | Overall Error (mm) | Yaw Error (°) | Pitch Error (°) | Roll Error (°) |
| --- | --- | --- | --- | --- |
| Overall | 1.05 (2.81) | 1.92 (7.25) | 2.35 (10.52) | 2.13(7.48) |
| 1920x1080px | 1.01 (2.95) | 1.57 (5.16) | 2.57 (9.30) | 2.04 (7.69) |
| 1640x1280px | 1.15 (2.57) | 2.34 (8.53) | 2.17(11.53) | 2.33(7.46) |

Finally, we validated the species classification and trajectory tracking performance in 3D-SOCS. We found that the YOLO model fine-tuned with great tit and blue tit annotations were highly accurate and largely outperformed out of the box methods based on pre-trained YOLO and zero-shot BioCLIP (Table S9). Tracking performance was high, with 90.0% of overall tracks with no mis-identification of IDs.

### 3.2 Estimating visual field use

Our stimulus display experiment provided a case study to use the capabilities of 3D-SOCS to determine the visual field use of *N* = 57 great tits. Each bird was recorded as visiting 3D-SOCS on average 18 times (range: 1-62). The orienting responses to the visual stimulus were defined as moments when the bird’s head was stationary (head-fixation); frames in which the bird exhibited rapid saccade-like head movements were excluded from the analysis.

We took several complementary approaches to assessing birds’ visual field use. Figure 2A summarizes head azimuth and elevation angles relative to all stimuli using a heatmap (see Figure S7 for each stimulus separately). Usage of both lateral and binocular fields are clearly shown by the high density regions. The hot-spots in the lateral field indicate the most likely positions of the optic axis (Itahara et al. 2022; Kano et al. 2022). When binning the marginal distributions of azimuth and elevation into 5° bins, the most observations fell between ±60° and ±65° azimuth and between −5° and −10° elevation, indicating the approximate locations of foveal projections.

**Figure 2:**
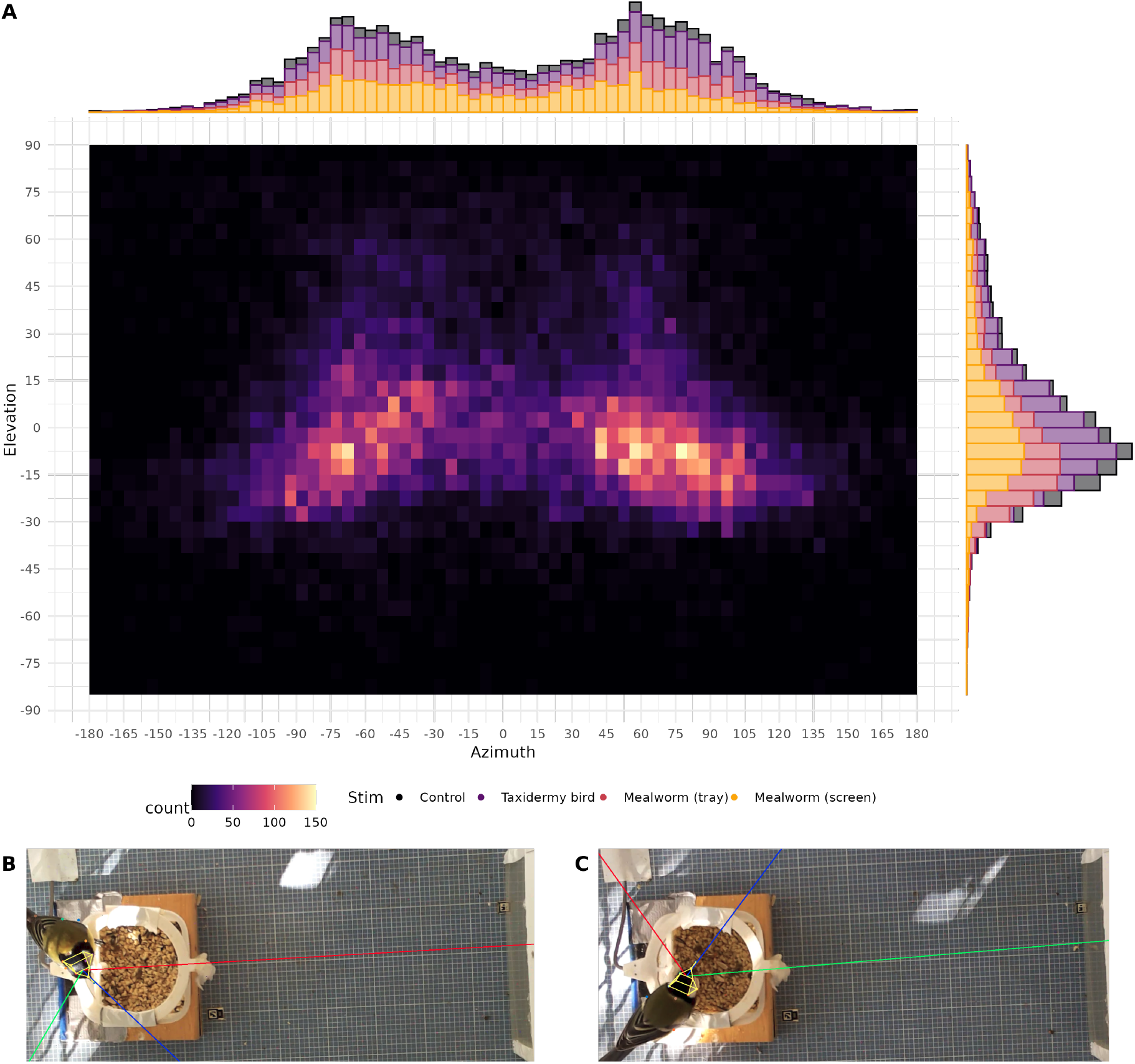
Summary of great tits’ visual field use. A) A heat-map illustrating the count of observations of head azimuth (x-axis, degrees) and elevation (y-axis, degrees) for all four stimuli. Histograms are included as marginal plots. Birds particularly used areas around ± 60° azimuth and −5° elevation to regard the stimuli. These areas likely encompasses their optic axes, as illustrated by frames taken when birds are regarding the stimulus with their **B)** left eye (red line rendered at −60° azimuth) and **C)** right eye (green line rendered at 60° azimuth).

In order to statistically validate this estimate, we used a Bayesian mixture model that estimated a 2dimensional normal distribution for each eye, composed of an azimuth and an elevation component, and a covariance matrix between azimuth and elevation parameters. A mixture parameter estimated the proportional contribution of each eye’s distribution to the overall distribution, although we note that this could not account for variation in eye preferences between individuals. Model results are reported in Table 3A, and fully summarized in Table S10. Visual field use was estimated to be most variable under the control condition, where no specific stimulus was present to observe. Across stimuli, we found a range of azimuth estimates between −65° to −56° for the left eye and 53° to 74° for the right eye. While mean estimates varied between stimuli, the mean absolute value of azimuth estimates was 59.58°, supporting our initial assessment of the optic axis at approximately 60°. The slightly wider right eye azimuth estimate for the taxidermy bird stimulus was either caused by 1) birds attending to the wooden stand the bird was attached to, rather than the bird itself, or 2) eye movement as birds held their heads more toward the direction of the exit for the taxidermy bird. The elevation was estimated to be higher than our initial assessment, although this was because elevation was modeled as a normal distribution, yet its marginal distribution was positively skewed. In light of the rotation error measured in section 3.1, we conclude the optic axes’ azimuth is approximately 60 ± 10° and elevation −5 ± 10°.

**Table 3:**
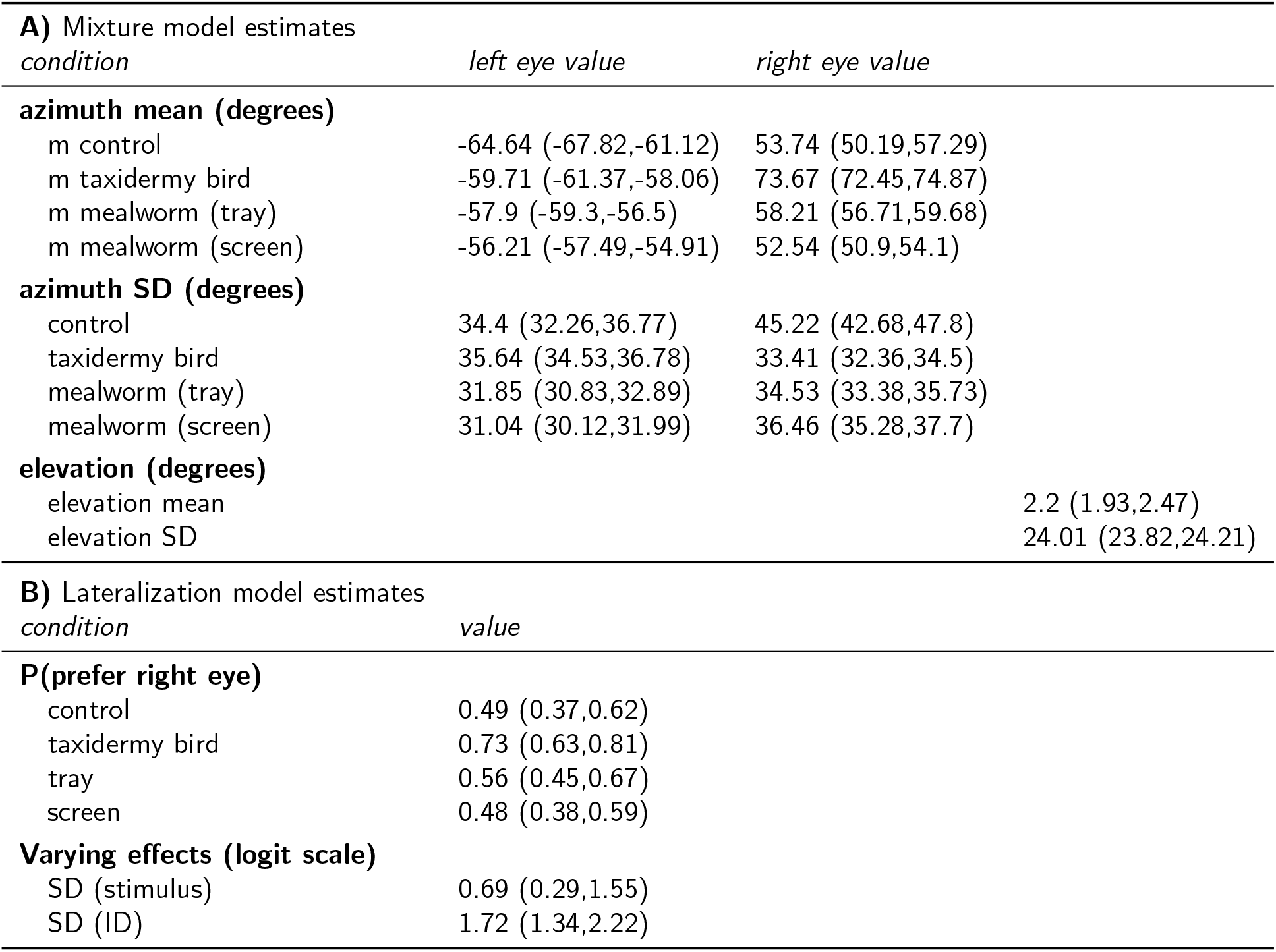
Summary of model estimates. Mean and 95% HPDI presented for all parameters. Models fit to data recorded within 3s after stimulus was presented. **A)** Bayesian mixture model described the estimated location of optic axes. **B)** Bayesian logistic regression with varying effects for stimulus type and bird identity was used to estimate differences in preferences for using left and right eyes when regarding stimuli.

In order to analyze bird’s preferences for using either eye, we fit a binomial model where the use of the lateral field of the right eye versus that of the left eye was predicted by varying effects for stimulus type and individual identity (Table 3B, fully summarized in Table S11). Age and sex showed no effect on lateralization, and these were excluded from the final model. We found more variation between birds than between stimuli (individual SD and 95% HPDI: 1.71(1.33, 2.2), stimuli: 0.81(0.31, 1.82). Interestingly, rather than a bimodal distribution of preferences, birds fell along a spectrum of lateralization (Figure S8). When controlling for differences between individuals, the taxidermy bird was estimated to elicit the strongest preference for the right eye (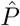 (use right lateral field) = .75). Weaker preference for the right eye was found in the control and food stimuli conditions, although the HPDI of these estimated probabilities crossed .5 indicating no strong preference.

Finally, to examine the temporal changes of birds’ orienting responses, we plotted the density of head angles from the moment of stimulus presentation as heat maps. To do this, we subset the data to observations with head elevation between -15 and 5 degrees (informed by the mixture model and system accuracy tests) to exclude observations where birds were likely not attending to the stimulus. This analysis provided a more nuanced picture of the birds’ visual behaviour over the time spent on the feeder (Figure 3). The clicking noise in the control condition appeared to elicit orienting responses, although this response was less structured compared to other stimuli and failed to maintain the birds’ attention for as long. When shown real mealworm stimuli, birds were more likely to use their binocular fields. In contrast, they primarily used their right eye to regard both the mealworm video stimulus and the taxidermy bird stimulus. They also regarded the taxidermy bird at a slightly wider angle.

**Figure 3:**
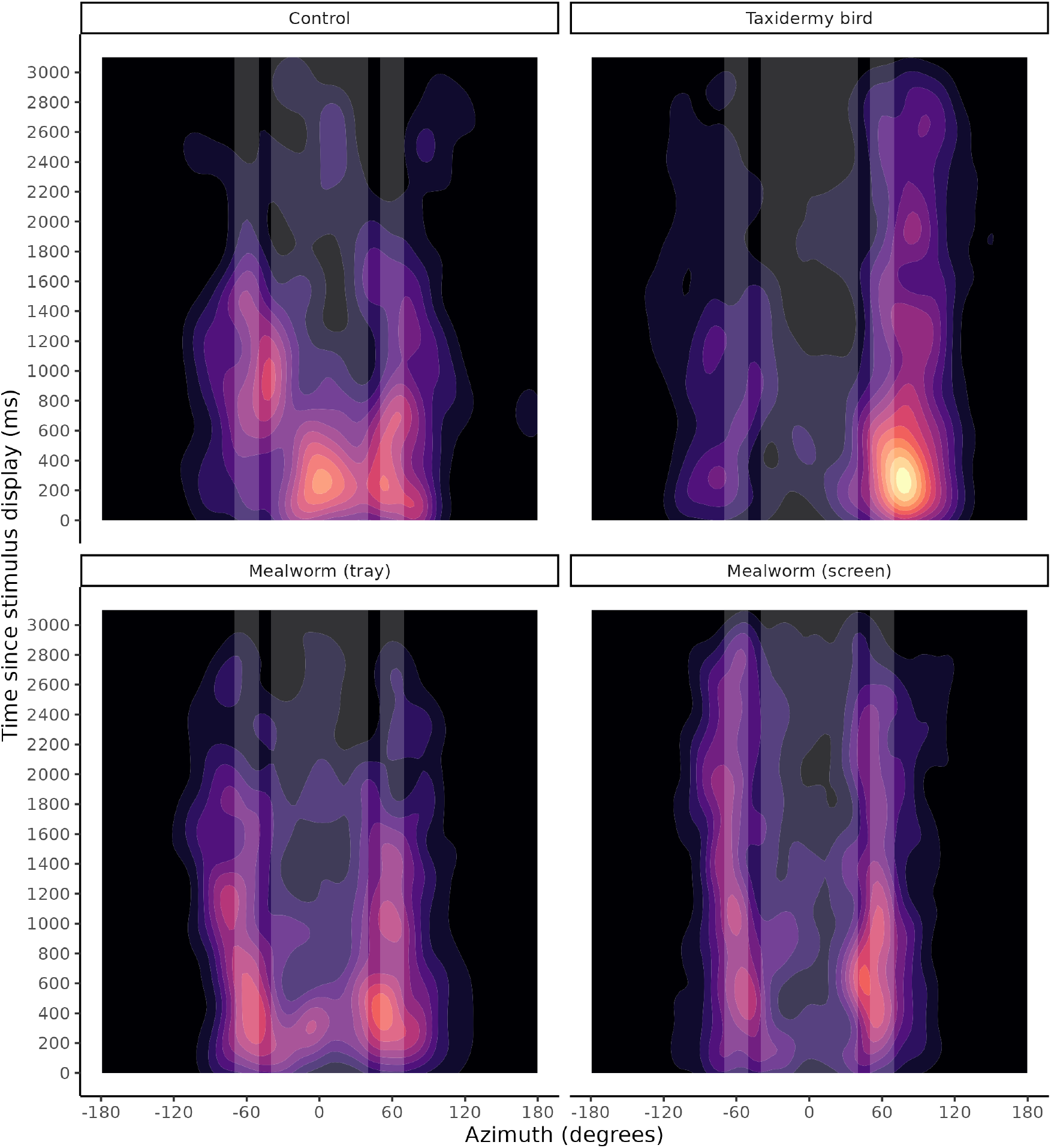
Gaze over time. Density of head azimuth over the 3s after the stimulus was displayed. Informed by our mixture model, data was subset to head elevations between -15 and 5 degrees to exclude probable non-looking observations. White rectangles indicate estimated optic axes from our mixture model, and likely binocular field based on estimates from related species (see discussion).

### 3.3 Population Monitoring

Next, we illustrate how 3D-SOCS can be used as a non-invasive population monitoring tool that provides insight into the distribution of weights of birds in a population without having to catch and weigh wild birds. We extracted the median convex hull surface area of both blue tits and great tits whose sex was known, and found a strong correlation between this measure and their mass measured during ringing (Pearson’s *r*[95%*CI*] = 0.900[0.843, 0.937], *t* = 16.888, df = 67, *p <* .001; Figure 4A). A simple model without covariates, simulating a researcher who might not have any data on individuals, found that surface area significantly predicted weight (Table S12; Figure S9). Model selection according to WAIC scores found the best fitting model accounted for varying intercepts for species, age and sex (Table S13 for model summary). Median body surface area as estimated from 3D-SOCS significantly predicted body weight from manually weighing undertaken when the birds were ringed (scaled surface area 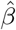 (95% HPDI) = 0.2(0.02, 0.38), Figure 4B). The contrast between varying intercepts for species also showed that the body size can significantly distinguish between great tit and blue tits 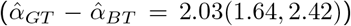. This reflects the known distribution of weights in these two species, where great tits are heavier than blue tits, with little to no overlap (mean weight GT=17.9g, BT=11.6g). However, we found no significant relationship between time-of-day or experimental day and variation in body size.

**Figure 4:**
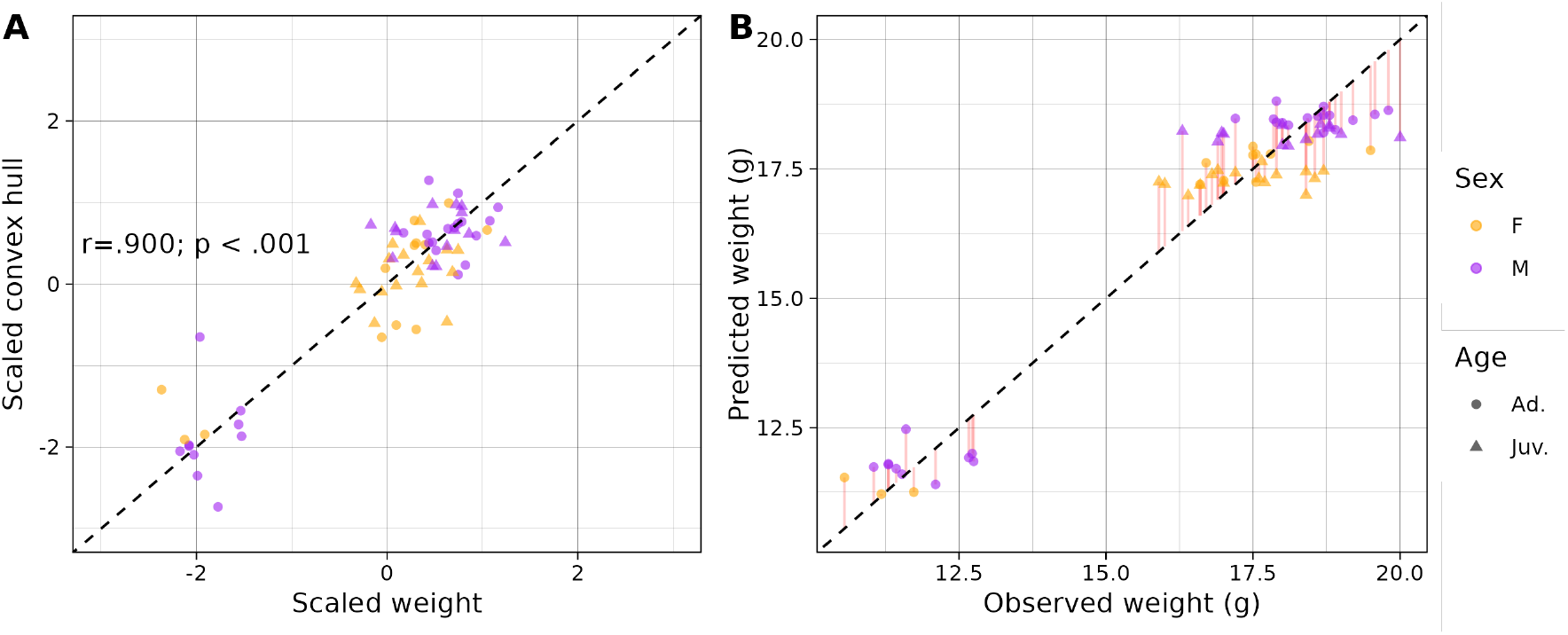
Median convex hull surface area predicts body mass. A) Correlation plot between scaled weight (x-axis) and scaled convex hull (y-axis), with Pearson’s *r* annotated. B) Actual weight (x-axis) versus predicted weight (y-axis) by best fitting model. Individuals’ sex and age are shown by color and shape, respectively. Residuals are shown in red. The bottom cluster is blue tits, the top cluster is great tits.

## 4 Discussion

We present 3D-SOCS as a flexible and inexpensive markerless posture tracking system for wild animals, and illustrated its capabilities with a common study system in behavioural ecology. To the best of our knowledge, this is the first system showcasing 3D markerless posture tracking of multiple animals from a wild population. Our evaluation of the system’s accuracy demonstrated that 3D-SOCS has no consistent systematic error in estimating location and angles of the head, and can recover keypoint estimates within 3mm of their true location. 3D-SOCS can also accurately classify great tit and blue tits at the species level, and track individuals reliably. Our experiment demonstrated the value of 3D tracking in examining great tits’ visual field use in response to visual stimuli. By tracking the birds’ head orientations, we were able to estimate the most frequently used visual field areas, primarily located near the optic axes, and also explored their lateralization of eye use. Finally, we show that body size estimation can be a useful proxy for species differentiation and body weight. We provide detailed documentation (available in supplementary text S1.1 and S1.2), and code (available in Section 5) for other researchers to deploy the system in the field who wish to quantify fine-scale behaviours in wild birds. Furthermore, 3D-SOCS is flexible in that any 2D keypoint detection model can be used rather than ours (e.g. SuperAnimalModel Ye et al. 2024), allowing it to potentially be used with other target species.

We validated 3D-SOCS performance by conducting a series of system accuracy tests by manually annotating multi-view frames, as well as comparing estimates to a marker-based motion capture system. Our evaluation demonstrated that the keypoint measurements in 3D-SOCS are normally distributed, with no systematic deviation from their ground truth values, and that errors typically fall within a 3mm range. By incrementally altering the number of cameras, we also showed that error rates are all comparable when reducing camera number, albeit with a trade-off in variability of the error and frame loss. Generally, more cameras produce less noisy and more consistent predictions, but a minimal setup of stereo cameras can still obtain accurate measures of 3D postures. Also importantly, 3D-SOCS is not limited to our setup with a metal cage. As long as two or more cameras can be mounted to provide multiple views of a scene, and a 2D posture detection model is available, 3D-SOCS can be effectively deployed across a variety of study systems and species.

We used 3D-SOCS to identify the visual field use of great tits with 3D tracking of head orientations. We found that the location of great tit’s optic axes are approximately *±*60° azimuth and *−*5° elevation. This result is largely consistent with previous literature in passerines. For example, captive zebra finches (*Taeniopygia guttata*) and crows (*Corvus macrorhynchos*) were previously found to have optic axes around ±60° (Bischof 1988; Itahara et al. 2024). For future researchers who would like to use 3D-SOCS to quantify the lateral gaze of great tits, we would suggest an uncertainty cone of 10° around our estimate of optic axes to account for both eye movement and rotational error. It was difficult to quantify the precise boundaries of the binocular field from our data, although birds certainly used their binocular field especially when they approached food items (Figures 2, 3). A prior study found that the binocular fields of tufted titmice and chickadees fell between ±26° azimuth when eyes were not rotated forward, and between ±40° when eyes were rotated forward (Moore et al. 2013). Given these species’ all fall in family Paridae, we speculate that tits’ values would be similar, although this would require empirical verification.

Furthermore, we observed lateralization in eye use in response to the different stimuli we presented. Hemispheric specialization could underlie the preferential use of either eye, although it has been argued that severe asymmetry should not be selectively favored (Vallortigara et al. 2001). Our results suggest that at the population level, tits were biased towards using their right eye (implicating the left hemispheric bias) especially when observing the taxidermy bird. In passerines, pigeons, and domestic chicks, the left hemisphere is generally superior in feature discrimination and memorization tasks (Prior 2006). It is therefore not surprising that great tits would also favor the right eye when attending to novel stimuli. However, this finding does contrast with the hypothesis that social cognition is dominated by the right hemisphere (Daisley et al. 2009; Salva et al. 2009), which is phylogenetically conserved (Hauser 1993; De Renzi et al. 1994; Sovrano et al. 1999; Kendrick 2006). However, quail (*Coturnix sp*.) were found to regard familiar individuals with their left eye, but to prefer the right eye for unknown conspecifics (Zucca et al. 2008). It may be that the tits follow a similar pattern, or that there was some uncertainty as to whether the stimuli represented a conspecific. This could be further explored by presenting a heterospecific stimulus.

We also demonstrated that body surface area can be used as a proxy for body weight, allowing for weights to be estimated using 3D-SOCS, providing valuable information about the overall individual variation in populations. However, our system is not well suited in measuring intra- or inter-day fluctuations because our definition of the convex hull of the body only included dorsal keypoints, while fat is deposited on the ventral side. Thus, if a customized system was designed for this purpose, measuring ventral keypoints, or using an electric scale would likely improve the estimation of body weights.

Although our illustrative experiments have highlighted the capabilities of 3D-SOCS, it is important to note that this system can be utilized across a variety of experimental paradigms. 3D-SOCS has the most potential for wild studies, especially when paired with cognitive tasks or experimental presentations, such as our stimulus presentation. However, 3D-SOCS could also allow for more naturalistic behaviour in laboratory experiments that measure gaze, which traditionally require restricting movement of animals (Tyrrell et al. 2014), or signalling, which can drastically change when animals are taken out of their natural context (Wilson 1992). Moreover, whether used in the wild or the lab, the tracked 3D postures can be utilized to automatically generate ethograms using either unsupervised or supervised algorithms (Dunn et al. 2021). The classified behaviours could then be used to quantify fine-scale variation in various interactions or displays such as fights or courtship displays at lekking sites (Janisch et al. 2021; Quesada et al. 2024). Finally, our finding that estimated body size serves as a reliable proxy for weight demonstrates a practical use-case for 3D-SOCS in quantifying body condition. This allows for repeated measurements without the need for catching, with various potential applications including monitoring morphological changes over time, such as in the observed link between climate change and decreasing body mass in great tits (Furness et al. 2019).

Computer vision methods for 3D posture estimation and automated behavioural classification have developed rapidly in recent years. However, these new methods often require idiosyncratic data (e.g. synchronized frames and calibration for 3D tracking). Yet, while more algorithms are being published, few studies focus on the deployment of these systems in experimental setups or in the field. As the field of computer vision in animals continue to grow, we encourage more 3D tracking systems designed to answer biologically meaningful questions related to behavioural ecology or conservation in wild-living animals. Innovations in system deployment is as important as designing novel algorithms and models. While 2D tracking has been widely used in the wild (Couzin et al. 2023), there is limited application of 3D tracking, as shown in Table 1. We hope 3D-SOCS sets a benchmark for future work, on how hardware and software can be developed in tandem and open-sourced for the scientific community to reproduce. However, there are still many potential improvements that could be made to 3D-SOCS. For example, it is limited to videos at 30 frames per second at a resolution of 1640×1232 px. By improving the spatial and temporal camera resolution, 3D-SOCS could be extended to study finer scale behaviours like signalling (Fusani et al. 2007) or wing beat frequencies (Steen 2014). Wireless camera synchronization could also be implemented to allow for easier deployment in the field. With the recent development and popularity of generalized posture models (Ye et al. 2024), models might soon require little to no training data, reducing the time needed for model training and validation.

In conclusion, we have demonstrated how 3D-SOCS offers a low-cost, non-invasive method for gaining unprecedented insights into the fine-scale detail of the behaviour of wild passerine birds. Our case study on the visual field of great tits suggests the possibility of inferring the area centralis in birds. Finally, we emphasize that 3D-SOCS is versatile in many respects: it can be deployed remotely in the field, it requires as few as two cameras, and with the appropriate inference models it could potentially be applied to a wide range of species.

## 5 Data Availability

Sample datasets for running the code and bounding box and keypoint annotations, as well as code and data for reproducing statistical analyses and main text figures are available via https://doi.org/10.5061/dryad. vq83bk429 (Chimento et al. 2025b). Code and instructions for creating and running 3D-SOCS yourself are available at https://github.com/alexhang212/3D-SOCS (Chimento et al. 2025a).

## 6 Acknowledgments

This work was supported by the Max Planck Society and the Centre for the Advanced Study of Collective Behaviour, funded by the Deutsche Forschungsgemeinschaft (DFG) under Germany’s Excellence Strategy (EXC 2117-422037984). Lucy M. Aplin and Michael Chimento were also partly funded by the Swiss State Secretariat for Education, Research and Innovation (SERI) under contract number MB22.00056. We thank Gustavo Alarcón-Nieto for helping with bird ringing, the members from the Comparative Sociality and Cognition Lab for the support and feedback, and MPI-AB veterinarians Inge Mueller and Daniel Zuñiga for their support.

## 7 Author Contributions

Conceptualisation - Michael Chimento, Alex Hoi Hang Chan, Lucy M. Aplin, Fumihiro Kano;

Methodology - Michael Chimento, Alex Hoi Hang Chan, Lucy M. Aplin, Fumihiro Kano;

Software - Michael Chimento, Alex Hoi Hang Chan;

Investigation - Michael Chimento, Alex Hoi Hang Chan;

Resources - Michael Chimento, Alex Hoi Hang Chan, Fumihiro Kano;

Writing Original Draft - Michael Chimento, Alex Hoi Hang Chan;

Writing Review & Editing - Michael Chimento, Alex Hoi Hang Chan, Lucy M. Aplin, Fumihiro Kano;

Visualization - Michael Chimento, Alex Hoi Hang Chan;

Supervision - Lucy M. Aplin, Fumihiro Kano;

Funding Acquisition - Michael Chimento, Lucy M. Aplin, Fumihiro Kano;

## References

Fisher, James and R. A. Hinde (1949). “The opening of milk bottles by birds”. In: British Birds 42, pp. 347– 357.

Gibb, John (1950). “The breeding biology of the Great and Blue titmice.” In: Ibis 92.4, pp. 507–539.

Perrins, Christopher M (1979). British tits. Vol. 62. HarperCollins.

Pratt, David W (1982). “Saccadic eye movements are coordinated with head movements in walking chickens”. In: Journal of Experimental Biology 97.1, pp. 217–223.

Bischof, Hans-Joachim (1988). “The visual field and visually guided behavior in the zebra finch (Taeniopygia guttata)”. In: Journal of Comparative Physiology A 163.3, pp. 329–337.

Wilson, Jeremy D (1992). “Correlates of agonistic display by great tits Parus major”. In: Behaviour, pp. 168– 214.

Hauser, Marc D (1993). “Right hemisphere dominance for the production of facial expression in monkeys”. In: Science 261.5120, pp. 475–477.

De Renzi, Ennio et al. (1994). “Prosopagnosia can be associated with damage confined to the right hemisphere—an MRI and PET study and a review of the literature”. In: Neuropsychologia 32.8, pp. 893– 902.

Lefebvre, Louis (1995). “The opening of milk bottles by birds: evidence for accelerating learning rates, but against the wave-of-advance model of cultural transmission”. In: Behavioural Processes 34.1, pp. 43–53.

Sovrano, Valeria Anna et al. (1999). “Roots of brain specializations: preferential left-eye use during mirror-image inspection in six species of teleost fish”. In: Behavioural brain research 106.1-2, pp. 175–180.

Vallortigara, Giorgio et al. (2001). “How birds use their eyes: opposite left-right specialization for the lateral and frontal visual hemifield in the domestic chick”. In: Current Biology 11.1, pp. 29–33.

Kendrick, Keith M (2006). “Brain asymmetries for face recognition and emotion control in sheep”. In: Cortex 42.1, pp. 96–98.

Prior, Helmut (2006). “Lateralization of spatial orientation in birds”. In: Behavioural and morphological asymmetries in vertebrates, pp. 75–85.

Fusani, Leonida et al. (2007). “High-speed video analysis reveals individual variability in the courtship displays of male golden-collared manakins”. In: Ethology 113.10, pp. 964–972.

Zucca, Paolo and Valeria A Sovrano (2008). “Animal lateralization and social recognition: quails use their left visual hemifield when approaching a companion and their right visual hemifield when approaching a stranger”. In: Cortex 44.1, pp. 13–20.

Daisley, Jonathan Niall et al. (2009). “Lateralization of social cognition in the domestic chicken (Gallus gallus)”. In: Philosophical Transactions of the Royal Society B: Biological Sciences 364.1519, pp. 965– 981.

Hawkes, Lucy A et al. (2009). “Climate change and marine turtles”. In: Endangered Species Research 7.2, pp. 137–154.

Salva, Orsola Rosa et al. (2009). “Lateralization of social learning in the domestic chick, Gallus gallus domesticus: learning to avoid”. In: Animal Behaviour 78.4, pp. 847–856.

Estók, Péter, Sándor Zsebők, and Björn M Siemers (2010). “Great tits search for, capture, kill and eat hibernating bats”. In: Biology letters 6.1, pp. 59–62.

Bonter, David N and Eli S Bridge (2011). “Applications of radio frequency identification (RFID) in ornithological research: a review”. In: Journal of Field Ornithology 82.1, pp. 1–10.

Morand-Ferron, Julie and John L Quinn (2011). “Larger groups of passerines are more efficient problem solvers in the wild”. In: Proceedings of the National Academy of Sciences 108.38, pp. 15898–15903.

Aplin, Lucy M et al. (2013). “Individual personalities predict social behaviour in wild networks of great tits (Parus major)”. In: Ecology letters 16.11, pp. 1365–1372.

Moore, Bret A et al. (2013). “Interspecific differences in the visual system and scanning behavior of three forest passerines that form heterospecific flocks”. In: Journal of Comparative Physiology A 199, pp. 263– 277.

Anderson, David J and Pietro Perona (2014). “Toward a science of computational ethology”. In: Neuron 84.1, pp. 18–31.

Farine, Damien R et al. (2014). “Collective decision making and social interaction rules in mixed-species flocks of songbirds”. In: Animal behaviour 95, pp. 173–182.

Lin, Tsung-Yi et al. (2014). “Microsoft COCO: Common Objects in Context”. In: Computer Vision – ECCV 2014. Ed. by David Fleet et al. Cham: Springer International Publishing, pp. 740–755. isbn: 978-3-319-10602-1.

Steen, Ronny (2014). “The use of a low cost high speed camera to monitor wingbeat frequency in hummingbirds (Trochilidae)”. In: Ardeola 61.1, pp. 111–120.

Tyrrell, Luke P et al. (2014). “A novel system for bi-ocular eye-tracking in vertebrates with laterally placed eyes”. In: Methods in Ecology and Evolution 5.10, pp. 1070–1077.

Aplin, Lucy M et al. (2015). “Experimentally induced innovations lead to persistent culture via conformity in wild birds”. In: Nature 518.7540, p. 538.

Farine, Damien R et al. (2015a). “Interspecific social networks promote information transmission in wild songbirds”. In: Proceedings of the Royal Society B: Biological Sciences 282.1803, p. 20142804.

Farine, Damien R et al. (2015b). “The role of social and ecological processes in structuring animal populations: a case study from automated tracking of wild birds”. In: Royal Society Open Science 2.4, p. 150057.

König, Barbara et al. (2015). “A system for automatic recording of social behavior in a free-living wild house mouse population”. In: Animal Biotelemetry 3, pp. 1–15.

Wilmers, Christopher C et al. (2015). “The golden age of bio-logging: How animal-borne sensors are advancing the frontiers of ecology”. In: Ecology 96.7, pp. 1741–1753.

Alarcón-Nieto, Gustavo et al. (2018). “An automated barcode tracking system for behavioural studies in birds”. In: Methods in Ecology and Evolution 9.6, pp. 1536–1547.

Butler, Shannon R, Jennifer J Templeton, and Esteban Fernández-Juricic (2018). “How do birds look at their world? A novel avian visual fixation strategy”. In: Behavioral ecology and sociobiology 72, pp. 1–11.

Flack, Andrea et al. (2018). “From local collective behavior to global migratory patterns in white storks”. In: Science 360.6391, pp. 911–914.

Iserbyt, Arne et al. (2018). “How to quantify animal activity from radio-frequency identification (RFID) recordings”. In: Ecology and Evolution 8.20, pp. 10166–10174.

Kano, Fumihiro et al. (2018). “Head-mounted sensors reveal visual attention of free-flying homing pigeons”. In: Journal of Experimental Biology 221.17.

Ling, Hangjian et al. (2018). “Simultaneous measurements of three-dimensional trajectories and wingbeat frequencies of birds in the field”. In: Journal of The Royal Society Interface 15.147, p. 20180653.

Mathis, Alexander et al. (2018). “DeepLabCut: markerless pose estimation of user-defined body parts with deep learning”. In: Nature neuroscience 21.9, pp. 1281–1289.

R Core Team (2018). R: A Language and Environment for Statistical Computing. R Foundation for Statistical Computing. Vienna, Austria. url: https://www.R-project.org/.

Dawson-Haggerty et al. (2019). trimesh. Version 3.2.0. url: https://trimesh.org/.

Furness, Euan N and Robert A Robinson (2019). “Long-term declines in winter body mass of tits throughout Britain and Ireland correlate with climate change”. In: Ecology and Evolution 9.3, pp. 1202–1210.

Graving, Jacob M et al. (2019). “DeepPoseKit, a software toolkit for fast and robust animal pose estimation using deep learning”. In: Elife 8, e47994.

Günel, Semih et al. (2019). “DeepFly3D, a deep learning-based approach for 3D limb and appendage tracking in tethered, adult Drosophila”. In: Elife 8, e48571.

Badger, Marc et al. (2020). “3D bird reconstruction: a dataset, model, and shape recovery from a single view”. In: European conference on computer vision. Springer, pp. 1–17.

Bala, Praneet C et al. (2020). “Openmonkeystudio: Automated markerless pose estimation in freely moving macaques”. In: BioRxiv, pp. 2020–01.

Beck, Kristina B, Damien R Farine, and Bart Kempenaers (2020). “Winter associations predict social and extra-pair mating patterns in a wild songbird”. In: Proceedings of the Royal Society B 287.1921, p. 20192606.

Fang, Cheng et al. (2020). “Comparative study on poultry target tracking algorithms based on a deep regression network”. In: Biosystems Engineering 190, pp. 176–183.

Ferreira André C et al. (2020). “Deep learning-based methods for individual recognition in small birds”. In: Methods in Ecology and Evolution 11.9, pp. 1072–1085.

Huang, Congzhentao et al. (2020). “End-to-end dynamic matching network for multi-view multi-person 3d pose estimation”. In: Computer Vision–ECCV 2020: 16th European Conference, Glasgow, UK, August 23–28, 2020, Proceedings, Part XXVIII 16. Springer, pp. 477–493.

Kane, Gary A et al. (2020). “Real-time, low-latency closed-loop feedback using markerless posture tracking”. In: Elife 9, e61909.

Mathis, Alexander et al. (2020). “A primer on motion capture with deep learning: principles, pitfalls, and perspectives”. In: Neuron 108.1, pp. 44–65.

Mathis, Mackenzie Weygandt and Alexander Mathis (2020). “Deep learning tools for the measurement of animal behavior in neuroscience”. In: Current opinion in neurobiology 60, pp. 1–11.

Chimento, Michael, Gustavo Alarcón-Nieto, and Lucy M Aplin (2021). “Population turnover facilitates cultural selection for efficiency in birds”. In: Current Biology 31.11, pp. 2477–2483.

Dunn, Timothy W et al. (2021). “Geometric deep learning enables 3D kinematic profiling across species and environments”. In: Nature methods 18.5, pp. 564–573.

Gosztolai, Adam et al. (2021). “LiftPose3D, a deep learning-based approach for transforming two-dimensional to three-dimensional poses in laboratory animals”. In: Nature methods 18.8, pp. 975–981.

Janisch, Judith et al. (2021). “Video recording and analysis of avian movements and behavior: insights from courtship case studies”. In: Integrative and Comparative Biology 61.4, pp. 1378–1393.

Jolles, Jolle W (2021). “Broad-scale applications of the Raspberry Pi: A review and guide for biologists”. In: Methods in Ecology and Evolution 12.9, pp. 1562–1579.

Joska, Daniel et al. (2021). “AcinoSet: a 3D pose estimation dataset and baseline models for Cheetahs in the wild”. In: 2021 IEEE international conference on robotics and automation (ICRA). IEEE, pp. 13901–13908.

Karashchuk, Pierre et al. (2021). “Anipose: a toolkit for robust markerless 3D pose estimation”. In: Cell reports 36.13.

Klarevas-Irby, James A, Martin Wikelski, and Damien R Farine (2021). “Efficient movement strategies mitigate the energetic cost of dispersal”. In: Ecology Letters 24.7, pp. 1432–1442.

Labuguen, Rollyn et al. (2021). “MacaquePose: a novel “in the wild” macaque monkey pose dataset for markerless motion capture”. In: Frontiers in behavioral neuroscience 14, p. 581154.

Lahoz-Monfort, José J and Michael JL Magrath (2021). “A comprehensive overview of technologies for species and habitat monitoring and conservation”. In: BioScience 71.10, pp. 1038–1062.

Maldonado-Chaparro, Adriana Alexandra, Wolfgang Forstmeier, and Damien R Farine (2021). “Relationship quality underpins pair bond formation and subsequent reproductive performance”. In: Animal Behaviour 182, pp. 43–58.

Mathis, Alexander et al. (2021). “Pretraining boosts out-of-domain robustness for pose estimation”. In: Proceedings of the IEEE/CVF winter conference on applications of computer vision, pp. 1859–1868.

Mitoyen Clémentine et al. (2021). “Female behaviour is differentially associated with specific components of multimodal courtship in ring doves”. In: Animal Behaviour 173, pp. 21–39.

Stan Development Team (2021). Stan Modeling Language Users Guide and Reference Manual. Version 2.27. url: http://mc-stan.org/.

Vehtari, Aki et al. (2021). “Rank-normalization, folding, and localization: An improved R for assessing convergence of MCMC (with Discussion)”. In: Bayesian analysis 16.2, pp. 667–718.

Walter, Tristan and Iain D Couzin (2021). “TRex, a fast multi-animal tracking system with markerless identification, and 2D estimation of posture and visual fields”. In: Elife 10, e64000.

Aharon, Nir, Roy Orfaig, and Ben-Zion Bobrovsky (2022). “BoT-SORT: Robust associations multi-pedestrian tracking”. In: arXiv preprint 2206.14651.

Cauchoix, Maxime et al. (2022). “The OpenFeeder: A flexible automated RFID feeder to measure interspecies and intraspecies differences in cognitive and behavioural performance in wild birds”. In: Methods in Ecology and Evolution 13.9, pp. 1955–1961.

Flack, Andrea et al. (2022). “New frontiers in bird migration research”. In: Current Biology 32.20, R1187–R1199.

Griebling, Hannah J et al. (2022). “How technology can advance the study of animal cognition in the wild”. In: Current Opinion in Behavioral Sciences 45, p. 101120.

Itahara, Akihiro and Fumihiro Kano (2022). ““Corvid Tracking Studio”: A custom-built motion capture system to track head movements of corvids.” In: Japanese Journal of Animal Psychology 72.1, pp. 1–16.

Kano, Fumihiro et al. (2022). “Head-tracking of freely-behaving pigeons in a motion-capture system reveals the selective use of visual field regions”. In: Scientific Reports 12.1, p. 19113.

Lauer, Jessy et al. (2022). “Multi-animal pose estimation, identification and tracking with DeepLabCut”. In: Nature Methods 19.4, pp. 496–504.

Pereira, Talmo D et al. (2022). “SLEAP: A deep learning system for multi-animal pose tracking”. In: Nature methods 19.4, pp. 486–495.

Tuia, Devis et al. (2022). “Perspectives in machine learning for wildlife conservation”. In: Nature communications 13.1, p. 792.

An, Liang et al. (2023). “Three-dimensional surface motion capture of multiple freely moving pigs using MAMMAL”. In: Nature Communications 14.1, p. 7727.

Chan, Alex Hoi Hang et al. (2023). “Comparison of manual, machine learning, and hybrid methods for video annotation to extract parental care data”. In: Journal of Avian Biology, e03167.

Chen, Jun et al. (2023). “Mammalnet: A large-scale video benchmark for mammal recognition and behavior understanding”. In: Proceedings of the IEEE/CVF conference on computer vision and pattern recognition, pp. 13052–13061.

Couzin, Iain D and Conor Heins (2023). “Emerging technologies for behavioral research in changing environments”. In: Trends in Ecology & Evolution 38.4, pp. 346–354.

Dunkley, Katie et al. (2023). “A low-cost, long-running, open-source stereo camera for tracking aquatic species and their behaviours”. In: Methods in Ecology and Evolution.

Ehlman, Sean M et al. (2023). “Leveraging big data to uncover the eco-evolutionary factors shaping behavioural development”. In: Proceedings of the Royal Society B 290.1992, p. 20222115.

Hu, Yujia et al. (Mar. 2023). “LabGym: Quantification of user-defined animal behaviors using learning-based holistic assessment”. In: Cell Reports Methods 3.3. issn: 2667-2375. doi: 10.1016/j.crmeth.2023.100415.

Jocher, Glenn, Ayush Chaurasia, and Jing Qiu (Jan. 2023). Ultralytics YOLO. Version 8.0.0. url: https://github.com/ultralytics/ultralytics.

Kings, Michael et al. (2023). “Wild jackdaws can selectively adjust their social associations while preserving valuable long-term relationships”. In: Nature Communications 14.1, p. 5103.

Koger, Benjamin et al. (2023). “Quantifying the movement, behaviour and environmental context of group-living animals using drones and computer vision”. In: Journal of Animal Ecology.

Luxem, Kevin et al. (2023). “Open-source tools for behavioral video analysis: Setup, methods, and best practices”. In: Elife 12, e79305.

Nagy, Máté et al. (2023). “SMART-BARN: Scalable multimodal arena for real-time tracking behavior of animals in large numbers”. In: Science Advances 9.35, eadf8068.

Naik, Hemal et al. (2023). “3D-POP-An automated annotation approach to facilitate markerless 2D-3D tracking of freely moving birds with marker-based motion capture”. In: Proceedings of the IEEE/CVF Conference on Computer Vision and Pattern Recognition, pp. 21274–21284.

Nourani, Elham et al. (2023). “Seabird morphology determines operational wind speeds, tolerable maxima, and responses to extremes”. In: Current Biology 33.6, pp. 1179–1184.

Schad, Lukas and Julia Fischer (2023). “Opportunities and risks in the use of drones for studying animal behaviour”. In: Methods in Ecology and Evolution 14.8, pp. 1864–1872.

Schofield, Daniel P et al. (2023). “Automated face recognition using deep neural networks produces robust primate social networks and sociality measures”. In: Methods in Ecology and Evolution 14.8, pp. 1937– 1951.

Wild, Sonja et al. (2023). “Manipulating actions: A selective two-option device for cognitive experiments in wild animals”. In: Journal of Animal Ecology 92.8, pp. 1509–1519.

Wild, Timm A et al. (2023). “Internet on animals: Wi-Fi-enabled devices provide a solution for big data transmission in biologging”. In: Methods in Ecology and Evolution 14.1, pp. 87–102.

Wiltshire, Charlotte et al. (2023). “DeepWild: Application of the pose estimation tool DeepLabCut for behaviour tracking in wild chimpanzees and bonobos”. In: Journal of Animal Ecology 92.8, pp. 1560–1574. doi: 10.1111/1365-2656.13932.

Xiao, Shiting et al. (2023). “Multi-view Tracking, Re-ID, and Social Network Analysis of a Flock of Visually Similar Birds in an Outdoor Aviary”. In: International Journal of Computer Vision 131.6, pp. 1532–1549.

Beck, Kristina B et al. (2024). “Experimental manipulation of population density in a wild bird alters social structure but not patch discovery rate”. In: Animal Behaviour 209, pp. 95–120.

Brookes, Otto et al. (Aug. 2024). “PanAf20K: A Large Video Dataset for Wild Ape Detection and Behaviour Recognition”. In: International Journal of Computer Vision 132.8, pp. 3086–3102. issn: 1573-1405. doi: 10.1007/s11263-024-02003-z.

Chan, Alex Hoi Hang et al. (2024). “YOLO-Behaviour: A simple, flexible framework to automatically quantify animal behaviours from videos”. In: bioRxiv. doi: 10.1101/2024.08.26.609387.

Chimento, Michael, Gustavo Alarcón-Nieto, and Lucy M Aplin (2024). “Immigrant birds learn from socially observed differences in payoffs when their environment changes”. In: PLoS biology 22.11, e3002699.

Delacoux, Mathilde and Fumihiro Kano (2024). “Fine-scale tracking reveals visual field use for predator detection and escape in collective foraging of pigeon flocks”. In: eLife 13.

Han, Yaning et al. (2024). “Multi-animal 3D social pose estimation, identification and behaviour embedding with a few-shot learning framework”. In: Nature Machine Intelligence, pp. 1–14.

Itahara, Akihiro and Fumihiro Kano (2024). “Gaze tracking of large-billed crows (Corvus macrorhynchos) in a motion capture system”. In: Journal of Experimental Biology, jeb–246514.

Kholiavchenko, Maksim et al. (Jan. 2024). “KABR: In-Situ Dataset for Kenyan Animal Behavior Recognition From Drone Videos”. In: Proceedings of the IEEE/CVF Winter Conference on Applications of Computer Vision (WACV) Workshops, pp. 31–40.

Quesada, Javier et al. (2024). “Recognizing interspecific dominance signals? Blue tits adjust nest defence based on great tit’s black bib size”. In: Ethology, e13460.

Raulo, Aura et al. (2024). “Social and environmental transmission spread different sets of gut microbes in wild mice”. In: Nature Ecology & Evolution, pp. 1–14.

Stevens, Samuel et al. (June 2024). “BioCLIP: A Vision Foundation Model for the Tree of Life”. In: Proceedings of the IEEE/CVF Conference on Computer Vision and Pattern Recognition (CVPR), pp. 19412– 19424.

Tkachenko, Maxim et al. (2024). Label Studio: Data labeling software. url: https://github.com/heartexlabs/label-studio.

Waldmann, Urs et al. (May 2024). “3D-MuPPET: 3D Multi-Pigeon Pose Estimation and Tracking”. In: International Journal of Computer Vision. issn: 1573-1405. doi: 10.1007/s11263-024-02074-y.

Ye, Shaokai et al. (2024). “SuperAnimal pretrained pose estimation models for behavioral analysis”. In: Nature Communications 15.1, p. 5165.

Chimento, Michael and Alex Hoi Hang Chan (2025a). alexhang212/3D-SOCS: 3D-SOCS release. Version v1.0. doi: 10.5281/zenodo.15146222. url: https://doi.org/10.5281/zenodo.15146222.

Chimento, Michael et al. (2025b). Data and code to reproduce “3D-SOCS: synchronized video capture for posture estimation”. url: 10.5061/dryad.vq83bk429.

