## Supplementary Materials for "Peering into the world of wild passerines with 3D-SOCS: synchronized video capture and posture estimation"

### Supplementary Text, Figures and Tables: Peering into the world of wild passerines with 3D-SOCS: synchronized video capture and posture estimation

immediate

April 25, 2025

#### Supplementary text


- **Text S1.1**: Setting up 3D-SOCS
- **Text S1.2**: 3D tracking pipeline


#### Supplementary figures


- **Figure S1**: Hardware schematic of 3D-SOCS.
- **Figure S2**: Head coordinate system.
- **Figure S3**: Saccade Statistics
- **Figure S4**: System accuracy and variance compared to data manually annotated in the wild.
- **Figure S5**: System accuracy validation histograms, six cameras.
- **Figure S6**: System accuracy validation histograms, stereo cameras.
- **Figure S7**: Visual field usage by stimulus.
- **Figure S8**: Individual variation in lateralization.
- **Figure S9**: Observed versus estimated weight from naive model
- **Figure S10**: System accuracy validation method.


#### Supplementary tables


- **Table S1**: Calibration drift over experimental days
- **Table S2**: Summary of parameters and priors for mixture model.
- **Table S3**: Summary table of error compared to multi-view manual annotations in the wild.
- **Table S4**: Tracking accuracy for full pipeline.
- **Table S5**: Accuracy, frame loss per post-processing step.
- **Table S6**: Detailed summary table of system accuracy tests in the SMART-BARN with six cameras.
- **Table S7**: Detailed summary table from system accuracy tests in the SMART-BARN with stereo cameras.
- **Table S8**: Summary table of system accuracy test in the SMART-BARN with differing number of cameras.
- **Table S9** Species classification performance of fine-tuned YOLO model compared to off-the-shelf models.

- **Table S10**: Model summary for azimuth elevation mixture model.
- **Table S11**: Model summary for lateralization.
- **Table S12**: Model summary for relationship between body size and weight for model without covariates.
- **Table S13**: Model summary for relationship between body size and weight for best fitting model.

#### Supplementary Videos

- Video S1: Qualitative results for 3D Tracking
- Video S2: Example of Great tit looking at stimulus
- Video S3: System accuracy test

### S1 Supplementary text

#### S1.1 Setting up 3D-SOCS

In the following sections we give critical instructions and details for successfully deploying 3D-SOCS. Running the system requires some proficiency in Linux and Python programming, and computer hardware and networking. An overview of the hardware setup is illustrated in Figure S1. pi0 is the "lead" CM4, and is responsible for acquiring the lead time from GPS satellites and signalling all other "follower" CM4s to start and stop recording. A list of required hardware and prices as of 2024 is provided in our github repository.

##### S1.1.1 Raspberry Pi Compute Module

Each camera is controlled by a Raspberry Pi Compute Module 4 (CM4) running the Raspberry Pi OS Linux distribution (`https://www.raspberrypi.com/software/`). These micro-computers are manufactured without any traditional I/O. Thus each CM4 must first be connected to an I/O board in order to be used (`https://www.raspberrypi.com/products/compute-module-4-io-board/`), which take a barrel-jack 12v power supply. Furthermore, CM4 come in a variety of options, depending on the amount of RAM required and the type of storage—eMMC (embedded NAND flash memory) versus "lite" (external microSD card). For our setup, we used a mixture of 2, 4 and 8GB RAM eMMC variants due to supply availability issues, but we suggest using the 8 GB RAM lite option. The lite option allows you to easily flash the operating system onto micro-SD cards using Raspberry Pi Imager (or equivalent) software. We recommend using 64GB SD cards so that onboard storage of videos is never a problem within a given day. As part of the supplementary data, we have provided full OS images that we used for both the "lead" CM4 and the "follower" CM4s. Once you have flashed the images to the microSD cards, attached the CM4 to the I/O boards, and inserted the microSD card, each CM4 can be operated like a normal desktop computer.

We attached the CM4 IO boards to a PVC plate using risers, and used plastic boxes with ventilation holes drilled into them as housing. CM4 have no built in heat management, and can easily cook themselves. We recommend purchasing heat sinks and fans to cool the system.

##### S1.1.2 GPS/GNSS time and PTP

One critical job that 3D-SOCS must do is synchronize the camera frame captures between all 6 follower CM4s. This requires all CM4 to synchronize their internal clocks as precisely as possible, which is accomplished using Precision Time Protocol (PTP), a network protocol for time synchronization (`https://standards.ieee.org/ieee/1588/6825/`. The lead CM4 acquires a lead time from GPS satellites using a Multi-constellation GPS / GNSS module and antenna manufactured by Timebeat (`https://www.timebeat.app/`). Follower CM4s then synchronize their own clocks to this lead time over their Ethernet connection via PTP. We recommend Timebeat's proprietary software to implement PTP, although any other PTP implementation could be used. If using the OS images we provided, be sure to update the license file, which can be obtained from Timebeat. Timebeat software will begin to run upon startup, and takes anywhere from 10 to 20 minutes to achieve sub-millisecond synchronization. GPS signal may not be found when operating 3D-SOCS inside of a building. For more details, please see Timebeat's documentation (`https://support.timebeat.app/hc/en-gb`).

##### S1.1.3 Cameras

Videos are recorded using Raspberry Pi V2 (overhead) and V3 (profile) cameras. The V3 camera's auto-focus mechanism is not suitable for extreme angles, and thus we used V2 cameras for overhead angles with a fixed focus. Cameras are connected to the CAM1 port on the CM4 IO boards. This port has a 22 pin connector, while the Raspberry Pi cameras have a larger 15 pin connector. This requires the use of a short adapter cable from 22 to 15 pin, an extender, and finally a 2m camera cable. Cameras must be connected before the CM4 is powered on to function.

On the lead CM4, the camera does not record video, but instead runs a motion-detection algorithm (figure S1). When motion is detected, a trigger signal (pin high) is sent via the GPIO5 pin (BCM naming convention) to follower CM4s who receive it on their GPIO5, ensuring that recording starts at the same moment, although due to variation cameras may start recording 1 or 2 frames apart. Videos are recorded for a fixed duration defined in the motion detection script, after which a second signal (pin low) is sent to stop recording.

Cameras are controlled using the Picamera2 python library (`https://datasheets.raspberrypi.com/camera/picamera2-manual.pdf`). We have created a custom encoder which uses a phase-locked loop algorithm that disciplines the frame rate of cameras to the system clock, which should be already synchronized via PTP. Camera frame rates are variable over time, and can quickly fall out of sync due to variation between cameras. The algorithm ensures that the encoder will continuously attempt to synchronize the frame captures throughout the recording. Properties of the cameras (i.e. resolution, focus of V3 cameras, white balance, etc.) can be manually set in the python script. For details related to focusing cameras, please see the readme document in the 3D-SOCS repository. Our script produces a metadata document for each video that can be used to process videos at a later step.

Before collecting data, the cameras must be calibrated. Calibration board throughout the arena, and ensuring a good number of detected points per camera for intrinsics and shared points for extrinsics. For intrinsic calibration, we first detect all available points from the calibration sequence, then used the camera calibration function from the opencv aruco library to obtain a calibration matrix and distortion coefficients for each camera. For extrinsic calibration, we use the stereocalibrate function for each camera pair, then transform all camera poses to be relative to one root camera, based on lowest reprojection error. Finally, we align and standardize the coordinate system across days by manually annotating and calculating the 3D positions of 6 distinct points in the scene with known distances, such that the origin (0,0,0) is right in front of the stimulus display device.

##### S1.1.4 Data management

Videos and associated metadata are written to the "Videos" directory of the CM4 with a filename that contains the camera name, date and a running index of the video number. The associated ".pts" file contained the metadata for each video. For each video frame we record the frame index, the system timestamp (taken from the system clock), the sensor timestamp (taken from camera clock), the time interval between the system timestamp of the current frame and last frame, the time interval between the sensor timestamp of the current frame and last frame, a synthetic time calculated from the start time of the video plus the cumulative sensor time, and a boolean indicating whether the frame was captured within 1ms of when it was supposed to be taken.

If you used 64 GB microSD cards, you should be able to comfortably record 1080p video for an entire day without having to move it onto external storage. Data management is largely the choice of the user. If desired, each CM4 could have its own external storage, and data could be directly written to this storage. Alternatively, the user could provide the lead CM4 with external storage, and use the supplied shell scripts to periodically move data onto this storage. Finally, the user could connect to the network switch with a laptop at the end of the day, and use the same script to pull videos onto their device. Once you have the data off of the system, it can then be fed into our 3D tracking pipeline detailed in the next section.

##### S1.1.5 Practical details

We strongly suggest to protect the electronics, as well as the camera cables, as wet cables introduce noise into the video signal. All electronics were housed in a plastic box which sat on top of the cage, protected by a roof fabricated from corrugated plastic and wood. We used two 10cm fans with passive heat-sinks to cool the Rasperry Pis. Keeping the system cool is critical, as overheated computers increase the frame-drop rate and make synchronization significantly worse. Camera cables were protected with 2cm diameter plumbing tubes.

With six cameras we could record for approximately 10 hours, with a 500W portable battery. Longer recording times would require a larger battery, and ideally larger capacity SD cards to avoid having to constantly pull data onto an external drive.

In this manuscript we used a computer with 64GB ram and a Nvidia RTX 3070 graphics card as a minimum requirement. We have not parallelized our processing scripts, so a CPU with any number of cores can be used. The processing speed is 0.94 frames per seconds (average across 3 repeated trials) on an Ubuntu machine with a 16GB Nvidia Geforce RTX 3070 GPU, 11th Gen Intel(R) Core(TM) i9- 11900 H @ 2.50GHz CPU, and Sandisk 2TB SSD.

#### S1.2 3D tracking pipeline

From the multi-view videos from 3D-SOCS, we first use YOLOv8 [1] to extract bounding boxes of the birds. The birds were then cropped using the bounding boxes and fed into a single animal Deeplabcut [2] model, to obtain 2D posture estimates. The 2D posture estimates were then smoothed using a rolling average filter with a window size of 3 frames to account for jittering of 2D point detections. With the detected 2D keypoints across all views, we then did 3D correspondence matching every frame, using an algorithm based on [3]. The algorithm iteratively matches points that are close in a 3D pose subspace, until a distance threshold is reached (set to 100mm in the current study). Next, the per-frame correspondence matching were mapped onto 2D tracks obtained from BoT-SORT, implemented using the YOLOv8 framework. To pair 2D tracks using the 3D correspondence information, we used a simple threshold by assuming a pair of 2D tracks as the same bird if the proportion of frames with the same correspondence is higher than 0.7. After the matching procedure, we get sets of 2D tracks that are assigned to the same individual, which was then used for triangulation with bundle adjustment using custom python scripts.

Once 3D posture estimates were obtained, we implemented a series of post-processing steps, as outlined in Table S5.

1) *Reprojection Filter*: 3D points were reprojected to each 2D camera, and if the 2D reprojection is outside any of the camera resolution, the point is filtered out.

2) *Distance filter*: each point was defined as the head or the body to compute a mean body position. Points with a distance of above 1 standard deviation above the mean were filtered out.

3) *Interpolation*: 3D points were interpolated using piecewise cubic hermite interpolating polynomial (PCHIP) interpolation, with a window size of 1. PCHIP was chosen over other interpolation options since it best preserves the rapid movement of the birds captured, without over-smoothing points. Interpolation were done two times, first after distance filter, second after object filter (Table S5).

4) *Object Filter*: Using measurements of each individual across the whole experimental period, we first constructed a unique median skeleton for each individual by transforming all head points into the local head coordinate system (see below). Using the median head object, we first calculated the rotation translation matrices of the detected head points for each given frame, then calculated the ideal head points in the global coordinate system by transforming the median skeleton based on the same rotation translation values. We then compared the ideal head points with the detected head points, then filtered out points with a difference of $> 50mm$ as outliers. The process is repeated until no outliers were detected, and the ideal points were taken as the final posture estimates. With this procedure, pairwise distances between each head point for each individual will always be identical across frames, improving consistency and minimizing frame loss.

#### S1.3 Defining the head coordinate system

We refer to Figure S2 for a detailed schematic on how the head coordinate system is computed. The head coordinate system is defined where the y-axis is a vector from the mid-point between the eyes towards the bill tip, and z axis as a vector normal perpendicular to the eye/bill tip plane. Finally, the x-axis is a vector perpendicular to the z-y plane. After the head coordinate system is defined, we computed the rotation translation matrices that transforms points from the world coordinate system to the head coordinate system, such that any subsequent detection from 3D-SOCS can then be transformed into the head coordinate system.

For a given object of interest (e.g stimulus), we first transform the object into the local head coordinate system, then calculate a modified spherical coordinate following [4–6]. In the modified spherical coordinates, azimuth and elevation of 0 represent the forward vector from midpoint of eyes towards the bill tip, with azimuth increasing as the object move towards the x-axis (right side of the head), and elevation increase as the object move towards the z-axis (above the eye/bill tip plane.) We refer to the code for detailed implementation of this procedure.

#### S1.4 System Accuracy Test

We used a taxidermy Great Tit and two aruco QR codes as detectable objects for the validation process. However, the SMART-BARN requires reflective markers to be placed in the points of interests in order to measure 3D coordinates, which would be a problem because placing a marker on detected keypoints (corners of a QR code or eye/beak of the bird) will affect the computer vision based detection methods of 3D-SOCS. To alleviate this problem, we mounted both the taxidermy bird and the QR code on a wooden plate, where we attached 4 reflective markers on. We then temporarily attached small circular reflective stickers on the points of interests (bill tip and eyes of the bird, four corners of the QR code, Figure S10), and recorded a short sequence from the motion capture system to establish the relative positions of the points of interests with the reflective markers. The reflective tape was then removed, and the 3D positions of the points of interests in the motion capture system can then be inferred each frame by calculating the rotation translation of the reflective markers per frame, using the object definition.

For the validation sequence, we first placed an ArUco QR code (6cm wide) in the middle of the volume, to act as the reference point. We then systematically rotated and moved the taxidermy great tit and another QR code within the volume for a total of around 20 minutes. We measured the detection error by calculating the distance of a given keypoint (e.g bill tip) relative to the reference QR code for both 3D-SOCS and SMART-BARN. The rotational error were then calculated by transforming the moving object to the object coordinate system of the reference QR code, then comparing the difference in rotation matrix between the two systems in terms of yaw pitch and roll. Since both systems are not strictly frame-synchronized, we slowly moved the taxidermy great tit through the volume and ensuring a long period of pause between each movement. We then used the SMART-BARN measurements to identify periods of inactivity (low rate of change), then took 30 frames of measurements in the mid-point of each of these periods of inactivity to ensure the error values were not influenced by system synchronization.

### S2 Supplementary figures

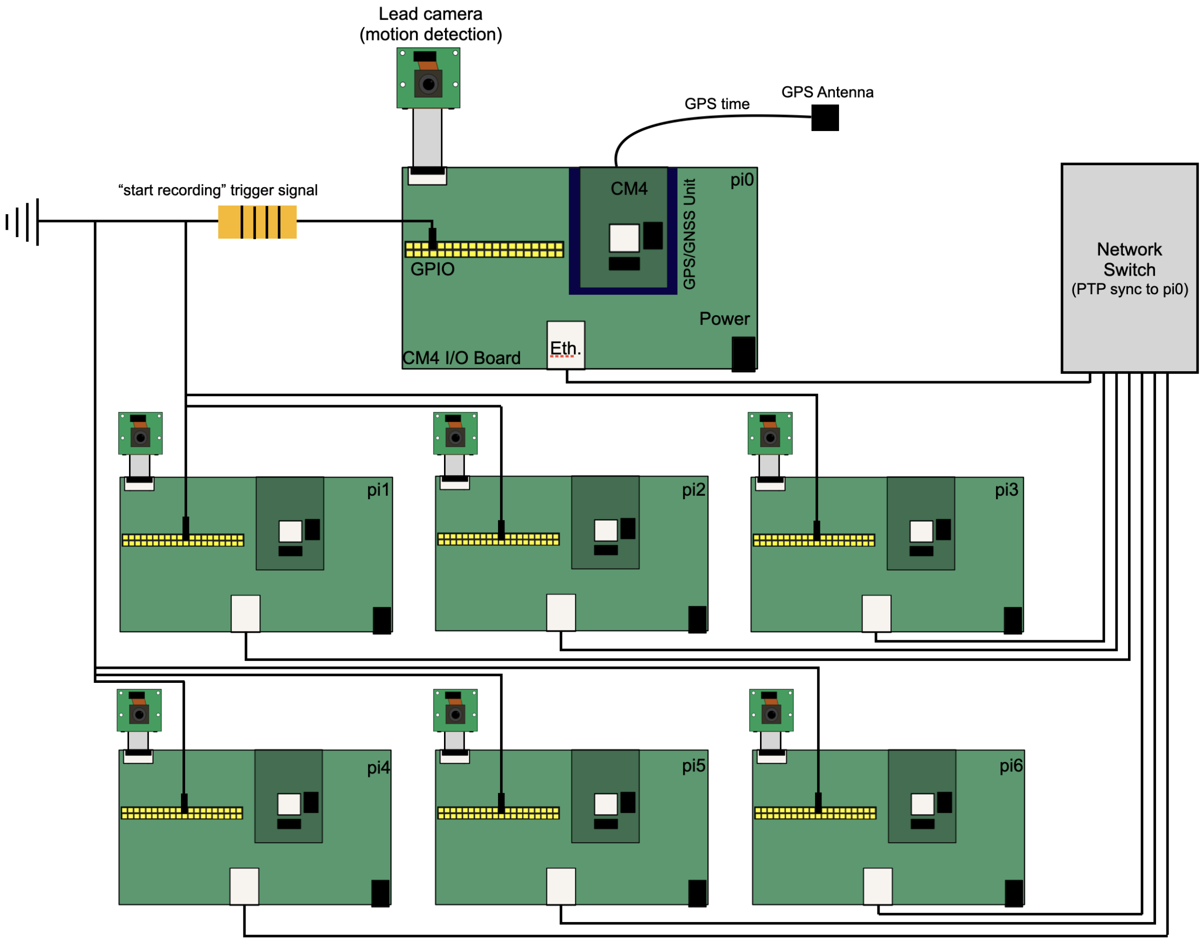


Figure S1: **Hardware schematic of 3D-SOCS**. One lead CM4 (pi0 in diagram) gets time from GNSS satellites, and also has a camera running motion detection algorithm. All 6 follower CM4 are networked to pi0 through a network switch, and synchronize their internal clocks to GNSS time. When motion is detected, pi0 sends an electrical signal over GPIO (with a resistor, importantly) for all follower CM4 to begin recording. Our customized camera encoder disciplines the frame rates of the cameras to GNSS time, ensuring highly synchronized frame capture. Video data can then be collected by connecting a laptop to the switch and downloading all data.

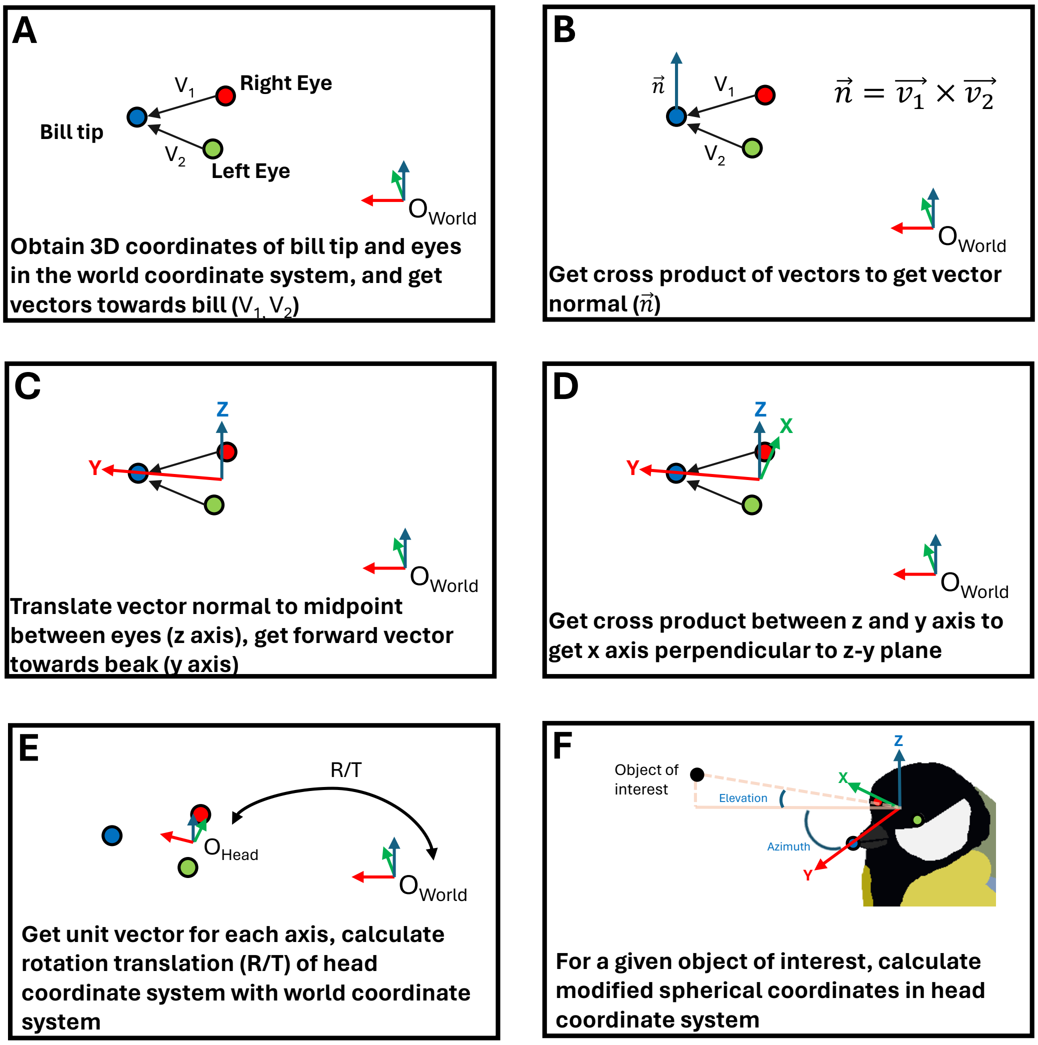


Figure S2: Head coordinate system. A) First obtain 3D coordinates of bill tip and eyes, then get vector from each eye towards the bill tip. B) get cross product between the vectors to get vector normal perpendicular to eye/bill tip plane. C) translate vector normal to be at the midpoint between eyes, defined as the origin of the new coordinate system. The vector normal is defined as the arbitrary z-axis, vector towards the bill tip is the y-axis. D) take the cross product of the z and y axis to get the x-axis, perpendicular to the z-y plane. E) get the unit vector of each axis and define as the head coordinate system, compute the rotation translation matrices that describes the new coordinate system. F) for a given object of interest in the head coordinate system, modified polar coordinates can be computed, where azimuth and elevation of 0 corresponds to the forward vector from eye mid-point to bill tip (i.e. y-axis).

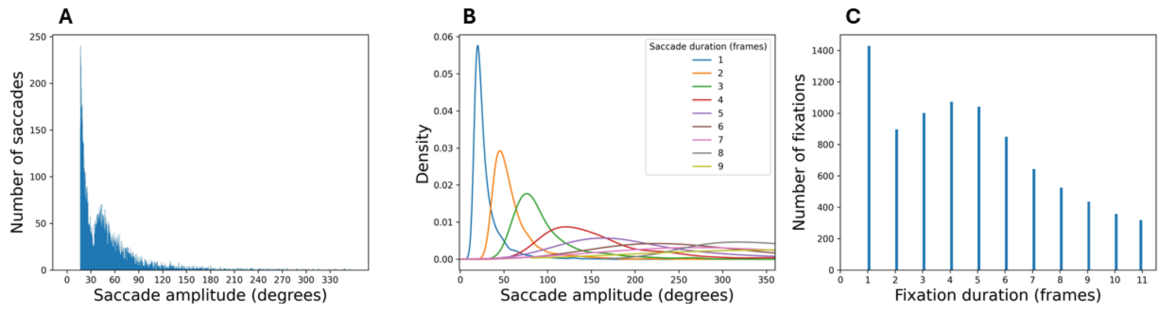


Figure S3: **Saccade Statistics**. Summary statistics of saccade-like head rotation movements in great tits, extracted by defining a saccade as changes of $> 500$ degrees per second (16.7 degrees per frame). A) Frequency of different saccade amplitudes. B) Density plot of saccade amplitudes grouped by the duration of the saccade. C) Frequency of the duration of fixations, defined as time between detected saccades

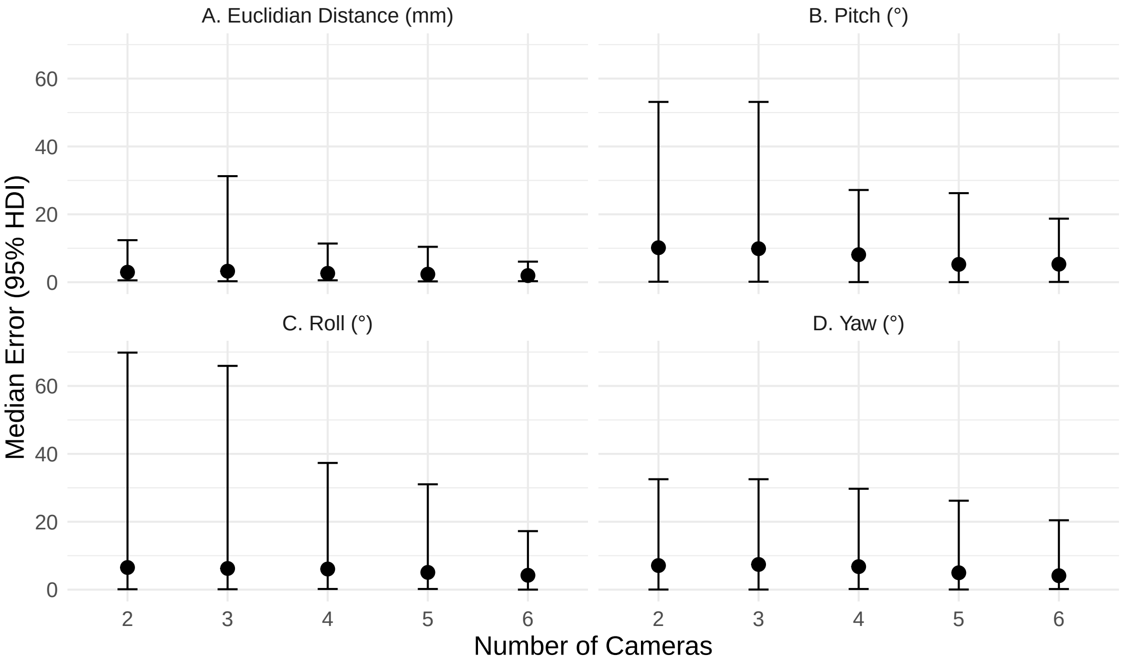


Figure S4: **Median error and 95% higher density interval (HDI) of 3D-SOCS in the wild**. Error values were generated by comparing detections against multi-view manual annotations of data collected by 3D-SOCS in the wild. A. Absolute euclidean distance, B-D, Pitch, Roll, Yaw respectively.

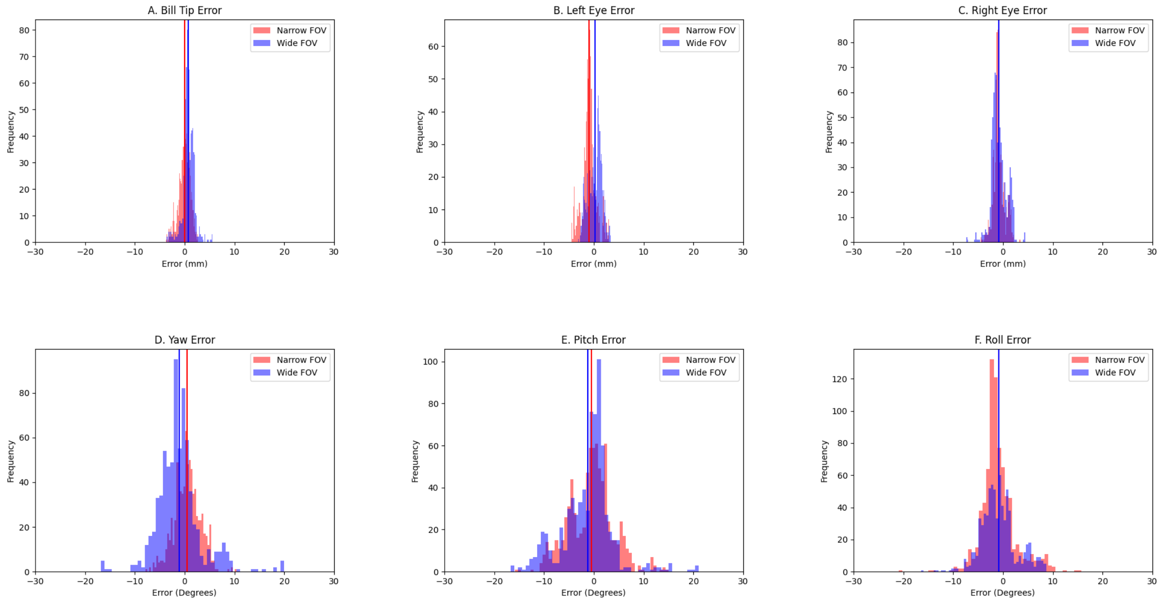


Figure S5: Histogram showing error distributions in the system accuracy test with taxidermy great tit, comparing 3D-SOCS with SMART-BARN. Red represents narrow field of view, blue represents wide field of view, with vertical lines representing the mean value. A-C) euclidean distance error between estimates. D-F) angular error between yaw, pitch and roll respectively.

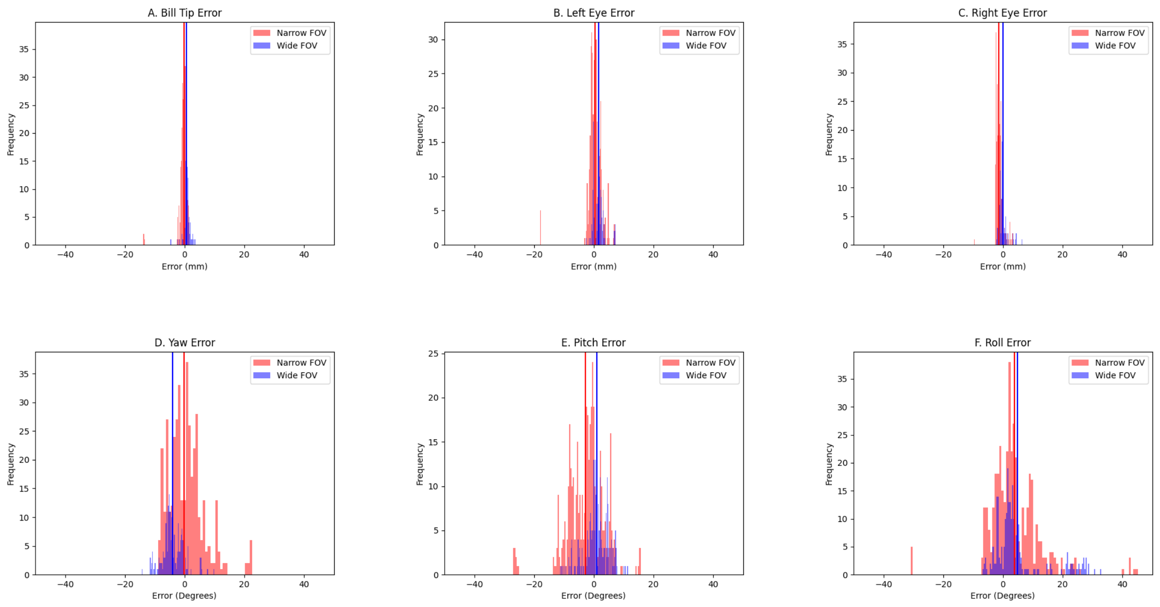


Figure S6: Histogram showing error distributions in the system accuracy test with taxidermy great tit, comparing 3D-SOCS with SMART-BARN with only 2 cameras. Red represents narrow field of view, blue represents wide field of view, with vertical lines representing the mean value. A-C) euclidean distance error between estimates. D-F) angular error between yaw, pitch and roll respectively.

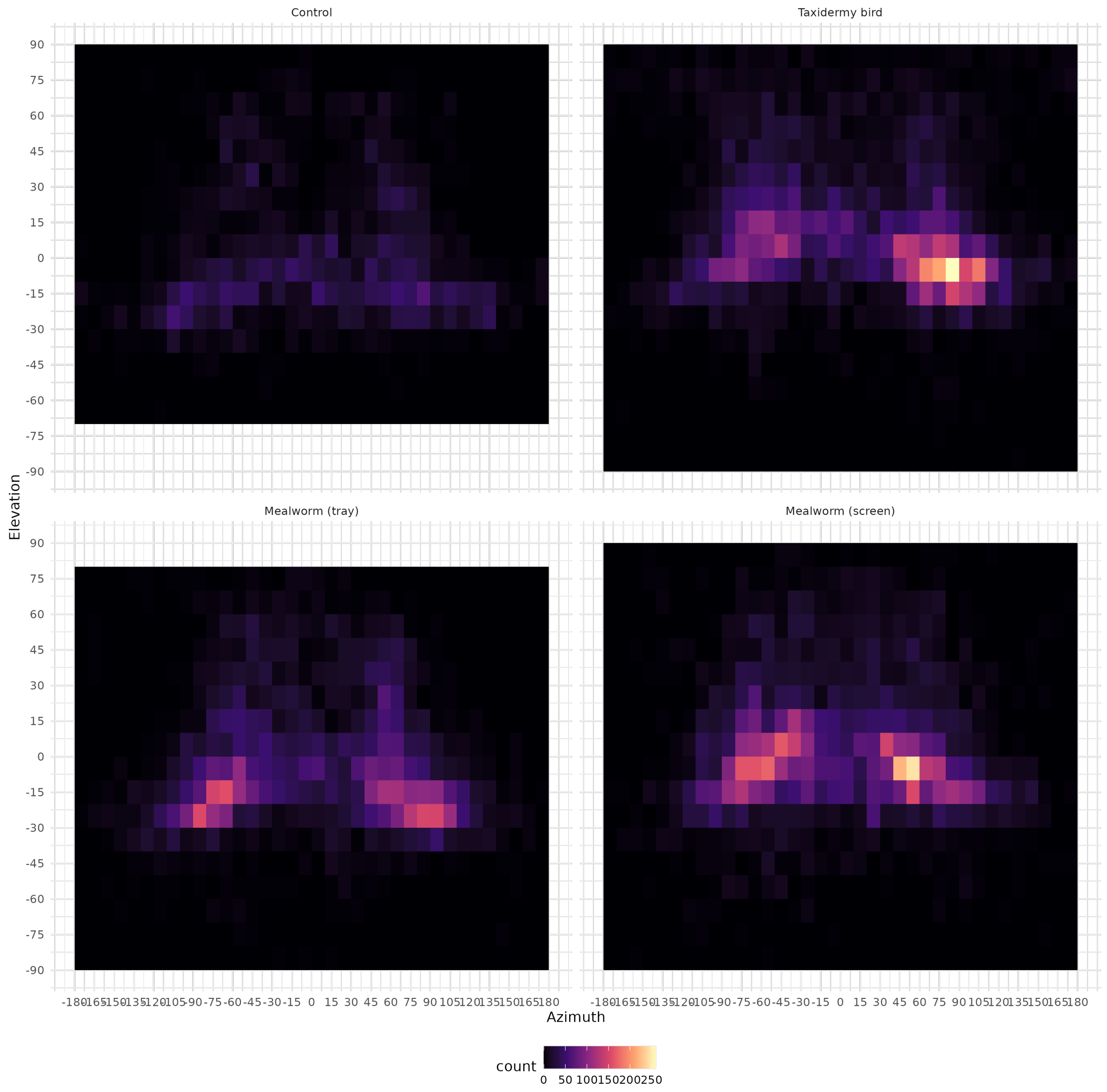


Figure S7: **Visual field usage by stimulus**. Heatmap of head azimuth (x-axis), head elevation (y-axis) of great tits relative to stimulus, recorded in the 3s following the stimulus display.

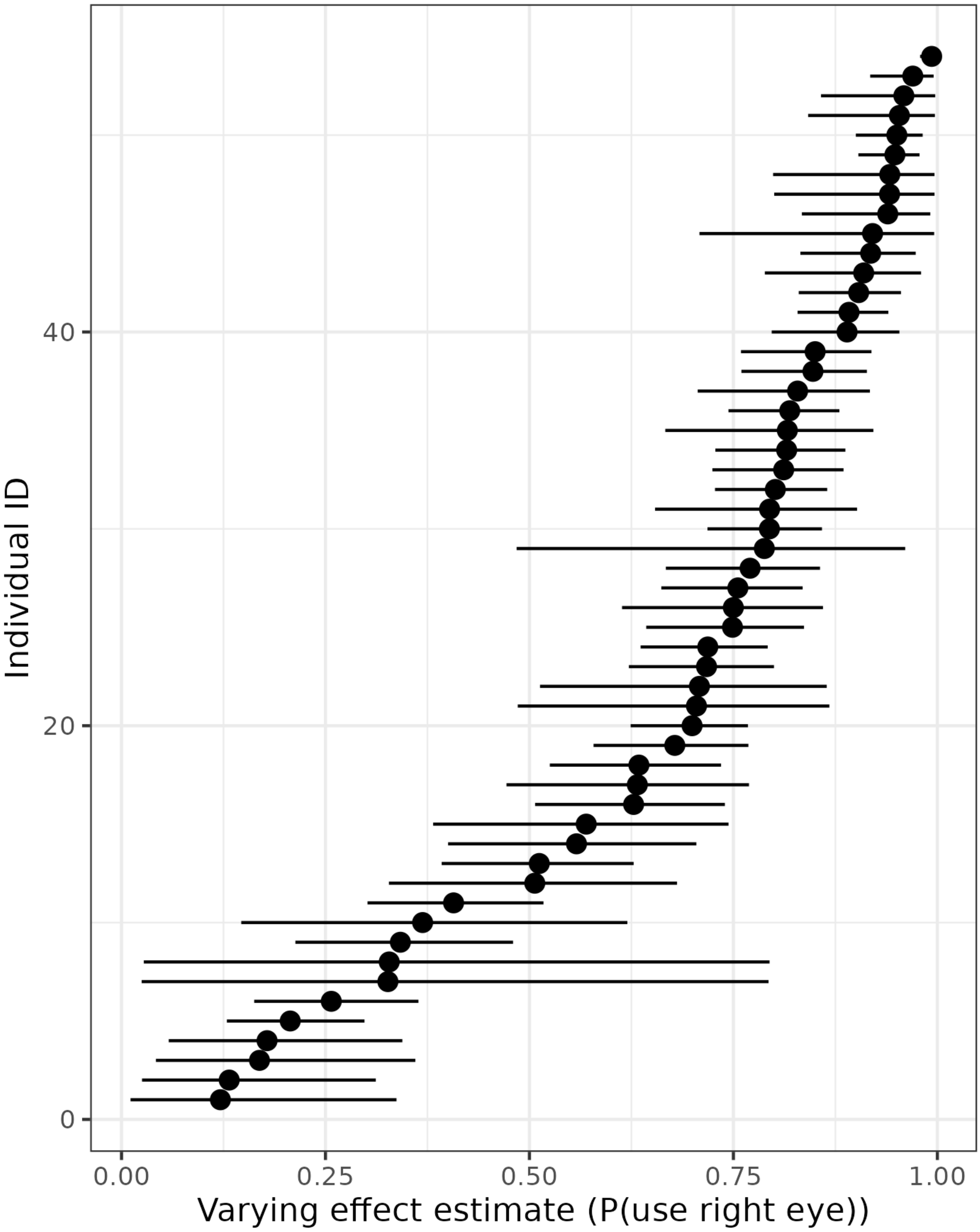


Figure S8: **Individual variation in lateralization**. Trellis plot of random effects from our lateralization model (transformed from logits to probabilities). Rather than a bimodal distribution of preferences, we found that birds fell along a spectrum of left/right eye preferences.

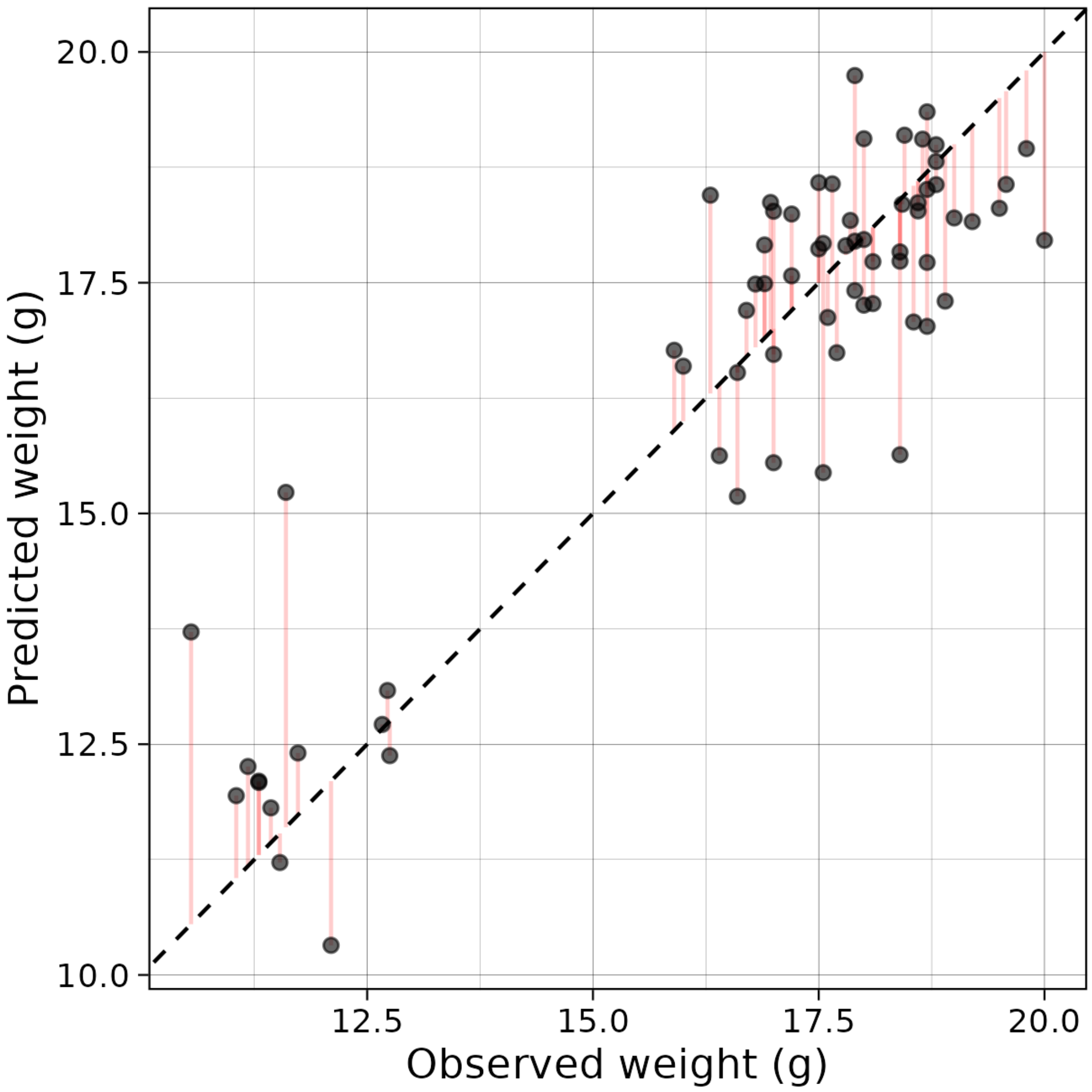


Figure S9: **Observed versus estimated weight from naive model**. This model included no information about age or sex, simulating a researcher that might not have these data, but still estimated a significant relationship between convex hull size and weight.

### S3 Supplementary tables

| Date | Mean Reprojection Error (px) | Mean 3D Error (mm) | Mean Reprojection Error (Day 1 calibration; px) | Mean 3D Error (Day 1 calibration; mm) |
|---|---|---|---|---|
| 05/11/2023 | 1.31 | 0.47 | 1.31 | 0.47 |
| 06/11/2023 | 1.83 | 0.46 | 5.19 | 0.61 |
| 07/11/2023 | 1.34 | 0.46 | 35.9 | 3.65 |
| 08/11/2023 | 1.28 | 0.43 | 28.5 | 2.75 |
| 09/11/2023 | 1.45 | 0.53 | 24.6 | 3.55 |
| 10/11/2023 | 1.53 | 0.41 | 24.0 | 3.62 |

Table S1: **Calibration drift over experimental days** Mean calibration error computed using charuco calibration board across experimental days, as mean reprojection error (px) and mean 3D error (mm) of distances between individual squares on the calibration board. Comparison made between using calibration parameters of the experimental day or the first experimental day (05/11/2023) to assess drift.

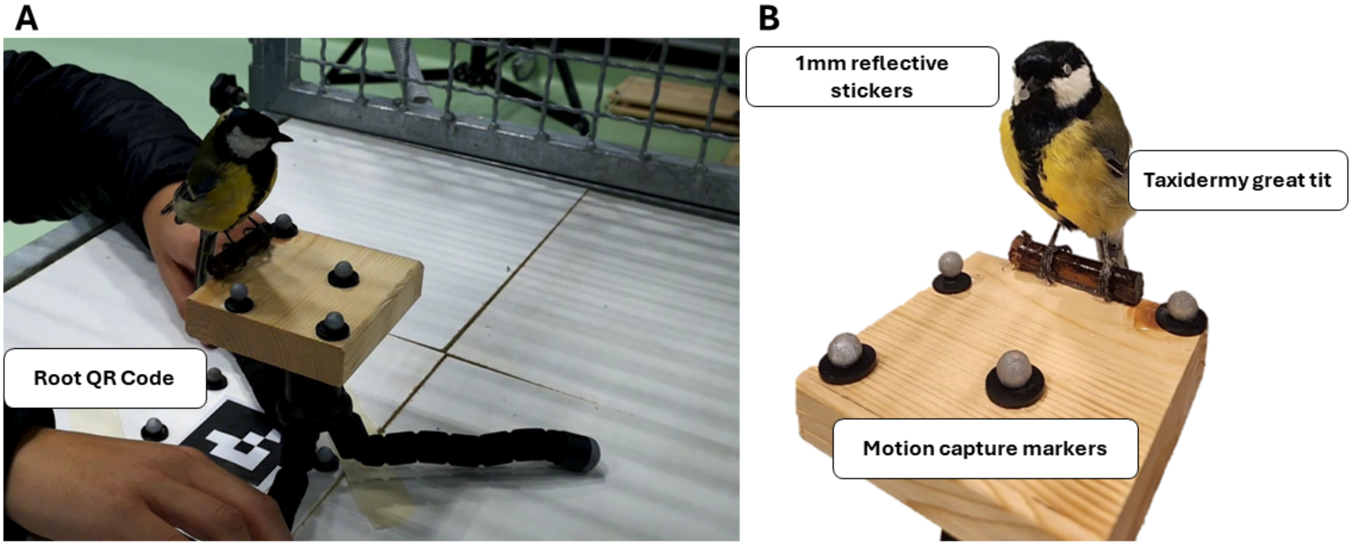


Figure S10: **System accuracy validation method.** A) a taxidermy great tit was placed within 3DSOCS and SMART-BARN, then moved and rotated systematically throughout the volume. The error is calculated by comparing the difference between the root QR code and keypoint of interest. B) 4 motion capture markers were attached to a wooden board, and 1mm reflective stickers were initially attached to the bill tip and eyes of the great tit to measure the distance between all points, then removed for validation trials.

| Parameter | Description | Prior Distribution |
| --- | --- | --- |
| $\theta_s$ | 2D simplex representing the mixing proportions for each of the $S$ stimuli. | Dirichlet(0.5, 2) |
| $\mu_{\text{az.},s}$ | ordered 2D-vector of locations for the azimuth component of the mixture, for each stimulus. | Normal(0, 1) |
| $\mu_{\text{elev.}}$ | Common location of mixture components for elevation | Normal(0, 1) |
| $\sigma_{\text{az.},s}$ | Scales of mixture components for azimuth, for each stimulus | Lognormal(0, 0.5) |
| $\sigma_{\text{elev.}}$ | Common scale of mixture components for elevation | Lognormal(0, 0.5) |

Table S2: Parameters and their prior distributions in the Bayesian hierarchical model.

| Number of Cameras | Overall Error (mm) | Yaw Error (°) | Pitch Error (°) | Roll Error (°) | Frame loss (%) |
|---|---|---|---|---|---|
| 2 | 2.95 (12.4) | 7.09 (32.5) | 10.2 (32.5) | 6.51 (69.8) | 58.4% |
| 3 | 3.25 (31.3) | 7.41 (32.5) | 9.88 (53.1) | 6.23 (65.9) | 48.6% |
| 4 | 2.62(11.1) | 6.78 (29.7) | 8.11 (27.2) | 6.06 (37.3) | 27.4% |
| 5 | 2.36 (10.4) | 4.97 (26.2) | 5.26 (26.2) | 5.08 (31.0) | 13.2% |
| 6 | 1.95 (6.07) | 4.10 (20.4) | 5.31 (18.7) | 4.22(17.2) | 12.3% |

Table S3: **Summary table of error compared to multi-view manual annotations in the wild** Median absolute error and upper 95% percentile interval in parenthesis, calculated as the overall accuracy of head points. Frame loss calulated as the percentage of frames with no measurements/ filtered out by the pipeline. Values presented for absolute euclidean distance, roll, pitch and yaw angles relative to ground truth. A value of zero would indicate a measurement that matched ground truth.

| Point Type | Median 2D detection error (px) | Median 3D Reprojection Error (px) | Average frame loss % |
|---|---|---|---|
| Overall | 0.24 | 4.56 | 18.7% |
| Head Points | 4.90 | 5.08 | 14.0% |
| Body Points | 3.26 | 3.53 | 27.3% |

Table S4: **Overall tracking performance statistics in 3DSOCS**. Overall median 2D detection error, median reprojection error and mean frame loss of detected keypoints in great tits captured by 3DSOCS. 2D detection error was extracted from the test set during network training. Frameloss was calculated by taking the mean frame loss across all keypoints.

| Post Processing Step | Median 3D Reprojection Error (px) | Average frame loss % |
|---|---|---|
| Triangulation | 3.06 | 30% |
| Reprojection Filter | 3.06 | 32% |
| Distance Filter | 3.01 | 42% |
| Interpolation 1 | 3.21 | 33% |
| Object Filter | 4.26 | 23% |
| Interpolation 2 (Final) | 4.56 | 19% |

Table S5: **Reprojection error and frame loss per post-processing step in the 3D tracking pipeline.** We report median 3D reprojection error and average frame loss at each step of the post-processing pipeline. We refer to the main text for more details.

| Resolution | Overall Error (mm) | Bill Tip Error (mm) | Left Eye Error(mm) | Right Eye Error(mm) |
|---|---|---|---|---|
| Overall | 1.05 (2.81) | 0.81 (2.45) | 1.07 (3.06) | 1.25 (2.72) |
| 1920x1080px | 1.00 (2.95) | 0.72 (2.38) | 1.06 (3.71) | 1.14 (2.57) |
| 1640x1280px | 1.15 (2.57) | 0.86 (2.73) | 1.09 (2.44) | 1.46 (2.95) |

Table S6: **Detailed summary table of system accuracy tests in the SMART-BARN with six cameras.** Median absolute error and upper 95% percentile interval in parenthesis. Values presented of head keypoints as well as roll, pitch and yaw angles relative to ground truth. A value of zero would indicate a measurement that matched ground truth. Keypoints estimated from six cameras, recorded at either a resolution of 1920x1080px (narrow FOV) or 1640x1280px (wide FOV).

| Resolution | Overall Error (mm) | Beak Error (mm) | Left Eye Error(mm) | Right Eye Error(mm) | Yaw Error (°) | Pitch Error (°) | Roll Error (°) |
|---|---|---|---|---|---|---|---|
| Overall | 0.90 (3.19) | 0.56 (2.28) | 1.12 (3.78) | 1.18 (3.11) | 4.05 (11.0) | 3.36 (11.4) | 3.39 (23.8) |
| 1920x1080px | 0.92 (2.68) | 0.49 (1.71) | 0.93 (4.82) | 1.49 (2.58) | 3.58 (11.9) | 4.07 (12.2) | 3.88 (19.9) |
| 1640x1280px | 0.87 (3.52) | 0.76 (2.86) | 1.61 (3.57) | 0.66 (4.06) | 4.77 (10.02) | 2.12 (7.68) | 2.98 (26.5) |

Table S7: **Summary table of system accuracy tests in the SMART-BARN with stereo cameras.** Median absolute error and upper 95% percentile interval in parenthesis. Values presented of head keypoints as well as roll, pitch and yaw angles relative to ground truth. A value of zero would indicate a measurement that matched ground truth. Keypoints estimated from only two cameras at once, recorded at either a resolution of 1920x1080px (narrow FOV) or 1640x1280px (wide FOV).

| Number of Cameras | Overall Error (mm) | Yaw Error (°) | Pitch Error (°) | Roll Error (°) |
|---|---|---|---|---|
| 2 | 0.90 (3.19) | 4.05 (11.0) | 3.36 (11.4) | 3.39 (23.8) |
| 3 | 1.22 (3.33) | 2.48 (9.32) | 3.87 (14.8) | 3.89 (16.1) |
| 4 | 1.11 (3.05) | 2.46 (8.31) | 2.31 (11.8) | 3.41 (13.7) |
| 5 | 1.08 (3.23) | 2.27 (9.69) | 2.53 (14.8) | 5.66 (9.98) |
| 6 | 1.05 (2.81) | 1.92 (7.25) | 2.35 (10.52) | 2.13(7.48) |

Table S8: **Summary table of system accuracy test in the SMART-BARN with differing number of cameras.** Median absolute error and upper 95% percentile interval in parenthesis, calculated as the overall accuracy of head points. Values presented for absolute euclidian distance, roll, pitch and yaw angles relative to ground truth. A value of zero would indicate a measurement that matched ground truth.

| Species/Model | Precision | Recall | f1-score | Accuracy |
|---|---|---|---|---|
| **A. Fine-tuned YOLO** | | | | |
| Great tit | 0.99 | 0.83 | 0.90 | |
| Blue tit | 0.93 | 0.87 | 0.90 | |
| Weighted Average | **0.97** | **0.84** | **0.90** | **0.84** |
| **B. Off the shelf YOLO + zero-shot bioclip** | | | | |
| Great tit | 0.80 | 0.17 | 0.28 | |
| Blue tit | 0.86 | 0.11 | 0.19 | |
| Weighted Average | 0.82 | 0.15 | 0.25 | 0.15 |
| **C. Ground truth bounding box + zero-shot bioclip** | | | | |
| Great tit | 0.79 | 0.61 | 0.69 | |
| Blue tit | 0.94 | 0.37 | 0.54 | |
| Weighted Average | 0.85 | 0.53 | 0.64 | 0.53 |

Table S9: **Species classification performance of fine-tuned YOLO model compared to off-the-shelf models.** Evaluation of species classification accuracy based on validation dataset used in YOLO training. Accuracy measures presented as: precision - proportion of predicted classes that were correct; recall - proportion of ground truth data that were correctly predicted; f1-score - combination metric of precision and recall; Accuracy - proportion of the full dataset that was correctly predicted. A) YOLO model used in the current study, fine-tuned for great tits and blue tits. B) Completely off the shelf pipeline, using pre-trained YOLOv8l to extract the "bird" class, then zero-shot bioclip for species identification. C) Using ground truth bounding box to crop images of birds, then zero-shot bioclip for species classification. Bold denotes best performing model.

| stimulus | value | eye | N_eff | Rhat |
|---|---|---|---|---|
| **theta (mixture)** | | | | |
| control | 0.39 (0.36,0.42) | left | 10555.58 | 1 |
| control | 0.61 (0.58,0.64) | right | 10555.58 | 1 |
| taxidermy bird | 0.44 (0.43,0.46) | left | 14767.15 | 1 |
| taxidermy bird | 0.56 (0.54,0.57) | right | 14767.15 | 1 |
| mealworm (tray) | 0.46 (0.44,0.47) | left | 15629.69 | 1 |
| mealworm (tray) | 0.54 (0.53,0.56) | right | 15629.69 | 1 |
| mealworm (screen) | 0.5 (0.49,0.52) | left | 13141.68 | 1 |
| mealworm (screen) | 0.5 (0.48,0.51) | right | 13141.68 | 1 |
| **azimuth mean (degrees)** | | | | |
| control | -64.64 (-67.82,-61.12) | left | 10192.88 | 1 |
| control | 53.74 (50.19,57.29) | right | 11229.35 | 1 |
| taxidermy bird | -59.71 (-61.37,-58.06) | left | 11710.05 | 1 |
| taxidermy bird | 73.67 (72.45,74.87) | right | 14959.71 | 1 |
| mealworm (tray) | -57.9 (-59.3,-56.5) | left | 13931.11 | 1 |
| mealworm (tray) | 58.21 (56.71,59.68) | right | 14056.76 | 1 |
| mealworm (screen) | -56.21 (-57.49,-54.91) | left | 13391.66 | 1 |
| mealworm (screen) | 52.54 (50.9,54.1) | right | 14390.78 | 1 |
| **azimuth SD (degrees)** | | | | |
| control | 34.4 (32.26,36.77) | left | 11715.79 | 1 |
| control | 45.22 (42.68,47.8) | right | 11676.26 | 1 |
| taxidermy bird | 35.64 (34.53,36.78) | left | 14639.81 | 1 |
| taxidermy bird | 33.41 (32.36,34.5) | right | 13264.03 | 1 |
| mealworm (tray) | 31.85 (30.83,32.89) | left | 14579.65 | 1 |
| mealworm (tray) | 34.53 (33.38,35.73) | right | 13810.12 | 1 |
| mealworm (screen) | 31.04 (30.12,31.99) | left | 14607.52 | 1 |
| mealworm (screen) | 36.46 (35.28,37.7) | right | 13421.95 | 1 |
| **elevation (degrees)** | | | | |
| elevation mean | 2.2 (1.93,2.47) | | 20948.83 | 1 |
| elevation SD | 24.01 (23.82,24.21) | | 22552.05 | 1 |
| **correlation btw. azimuth, elevation** | | | | |
| control | 20.86 (17.51,24.12) | left | 14946.64 | 1 |
| control | -15.45 (-18.36,-12.5) | right | 20587.70 | 1 |
| taxidermy bird | 5.43 (3.49,7.35) | left | 18009.11 | 1 |
| taxidermy bird | -14.98 (-16.74,-13.16) | right | 18392.60 | 1 |
| mealworm (tray) | 22.42 (20.61,24.17) | left | 21264.24 | 1 |
| mealworm (tray) | -22.19 (-23.86,-20.45) | right | 18918.45 | 1 |
| mealworm (screen) | 13.61 (11.7,15.5) | left | 20528.76 | 1 |
| mealworm (screen) | -17.57 (-19.36,-15.78) | right | 21292.67 | 1 |

Table S10: **Model summary for azimuth elevation mixture model.** Results for Gaussian mixture model which was used to statistically assess the location of the optic axes. Reported are parameter values (95% HPDI), which eye the estimate applies to, number of effective samples and Rhat value.

| parameter | value | P(prefer right eye) | n_eff | Rhat |
|---|---|---|---|---|
| beta (control) | -0.04 (-0.51,0.48) | 0.49 | 2234.89 | 1 |
| beta (taxidermy bird) | 0.97 (0.54,1.44) | 0.73 | 1680.07 | 1 |
| beta (tray) | 0.24 (-0.19,0.7) | 0.56 | 1698.36 | 1 |
| beta (screen) | -0.07 (-0.48,0.37) | 0.48 | 1559.10 | 1 |
| SD (stimulus) | 0.69 (0.29,1.55) | 0.67 | 7484.31 | 1 |
| SD (ID) | 1.72 (1.34,2.22) | 0.85 | 11016.03 | 1 |

Table S11: **Model summary for lateralization.** Results for logistic regression with varying effects for stimulus and ID, where the probability of using the right eye was the response variable. Reported are parameter values on the logit scale (95% HPDI), probability point estimate, number of effective samples and Rhat value.

| parameter | value | n_eff | Rhat |
|---|---|---|---|
| beta (scaled SA) | 0.9 (0.79,1) | 11361.16 | 1 |
| intercept | 0 (-0.11,0.11) | 11378.07 | 1 |
| SD | 0.45 (0.38,0.53) | 10750.51 | 1 |

Table S12: **Model summary for relationship between body surface area and weight without covariates.** Great tits' mean body weight (scaled) predicted by body SA (scaled, estimated from keypoints), and species class. Reported are parameter values (95% HPDI), number of effective samples and Rhat value.

| parameter | value | n_eff | Rhat |
|---|---|---|---|
| beta (scaled SA) | 0.2 (0.02,0.38) | 9866.70 | 1 |
| intercept | -0.28 (-1.79,1.24) | 7122.41 | 1 |
| varying intercept (female) | -0.24 (-1.49,0.99) | 6757.89 | 1 |
| varying intercept (male) | -0.01 (-1.26,1.2) | 6803.21 | 1 |
| varying intercept (adult) | -0.07 (-1.3,1.18) | 6702.43 | 1 |
| varying intercept (juvenile) | -0.18 (-1.4,1.05) | 6680.17 | 1 |
| varying intercept (blue tit) | -1.15 (-2.43,0.12) | 6614.23 | 1 |
| varying intercept (great tit) | 0.88 (-0.42,2.12) | 6713.33 | 1 |
| SD | 0.31 (0.26,0.37) | 10149.49 | 1 |

Table S13: **Model summary for relationship between body size and weight for best fitting model.** Great tits' mean body weight (scaled) predicted by body SA (scaled, estimated from keypoints), and species class. Reported are parameter values (95% HPDI), number of effective samples and Rhat value.

### S4 Supplementary Videos

#### S4.1 Supplementary Video S1

**Qualitative results for multi-individual 3D tracking in 3D-SOCS**. Left hand panels show detected 2D keypoints over six synchronized cameras. Detected 3D points were reprojected back to 2D and shown in the right panel, with line representing foveas, 60°left and right from the bill tip projection. Video was slowed down to 0.5x for visualization.

#### S4.2 Supplementary Video S2

**Example of great tit looking at screen mealworm stimulus**. Detected 3D points were reprojected back to 2D, with lines representing fovea projections, 60°left and right from bill tip

#### S4.3 Supplementary Video S3

**Example sequence in system accuracy test with taxidermy great tit**. A taxidermy great tit was systematically rotated in space, as part of the system accuracy test. The detected 3D keypoint estimate from 3D-SOCS is reprojected to 2D. The video was shown in 4x speed.

### References


1. Jocher, G., Chaurasia, A. & Qiu, J. *Ultralytics YOLO* version 8.0.0. Jan. 2023. `https://github.com/ultralytics/ultralytics`.
2. Mathis, A. *et al.* DeepLabCut: markerless pose estimation of user-defined body parts with deep learning. *Nature neuroscience* **21,** 1281–1289 (2018).
3. Huang, C. *et al. End-to-end dynamic matching network for multi-view multi-person 3d pose estimation* in *Computer Vision–ECCV 2020: 16th European Conference, Glasgow, UK, August 23–28, 2020, Proceedings, Part XXVIII 16* (2020), 477–493.
4. Kano, F., Naik, H., Keskin, G., Couzin, I. D. & Nagy, M. Head-tracking of freely-behaving pigeons in a motion-capture system reveals the selective use of visual field regions. *Scientific Reports* **12,** 19113 (2022).
5. Delacoux, M. & Kano, F. Fine-scale tracking reveals visual field use for predator detection and escape in collective foraging of pigeon flocks. *eLife* **13** (2024).
6. Itahara, A. & Kano, F. Gaze tracking of large-billed crows (Corvus macrorhynchos) in a motion capture system. *Journal of Experimental Biology,* jeb–246514 (2024).
